# Reconciling contradictory models of subthalamic nucleus contributions to basal ganglia beta oscillations

**DOI:** 10.64898/2026.01.26.701663

**Authors:** Ka Nap Tse, G. Bard Ermentrout, Jonathan E. Rubin

## Abstract

Recent computational studies of Parkinson’s disease have yielded contradictory findings regarding the role of the subthalamic nucleus (STN) in pathological beta oscillations, with some models implicating STN as essential for beta generation and others suggesting that STN suppresses oscillations. This work addresses these discrepancies by systematically investigating how the specific features of the integrate-and-fire neurons used in these models influence simulated basal ganglia network dynamics. Using both rate models and spiking network simulations incorporating coupled subthalamopallidal and pallidostriatal circuits, we demonstrate that the choice between leaky integrate-and-fire (LIF) and quadratic integrate-and-fire (QIF) models to represent STN neurons fundamentally impacts the phase relationship between STN and external globus pallidus prototypical (Proto) neuron populations. QIF STN neurons establish in-phase coupling with Proto neurons, which enhances beta oscillation amplitude, while LIF STN neurons develop anti-phase relationships, which suppresses beta power. Through intervention experiments and parameter sweeps across physiologically relevant firing rates, we show that these phase-related effects persist robustly across network conditions, and we mathematically establish conditions under which these results are guaranteed to hold. Our findings reveal that the fundamental mathematical structure underlying spike generation, rather than other biophysical details, determines whether the subthalamopallidal loop acts as a beta amplifier or suppressor. This mechanistic insight reconciles contradictory findings in the literature, demonstrates that seemingly minor modeling choices can have profound consequences for understanding disease mechanisms and therapeutic targets, and offers predictions for determining which model framework reflects the biological reality.

**Author summary:** Substantial work has explored the mechanisms underlying enhanced beta oscillations in the basal ganglia, motivated by their potential relevance to parkinsonian conditions and associated treatments. Often inferences about these mechanisms are based on simulations and reasoning that focus on features of network connectivity. We show that in fact the specific dynamical properties of the neurons in these circuits can strongly influence their emergent dynamics, with completely opposing effects arising in a given network structure depending on which neuron model is used, and we explain the factors underlying this divergence. Based on these factors, the determination of a small set of neuron properties in future biological experiments will lead to predictions about the mechanisms that can generate beta oscillations in the parkinsonian basal ganglia.

## Introduction

The basal ganglia network comprises interconnected subcortical nuclei including the striatum with its spiny projection neurons and fast-spiking interneurons [1–3], the subthalamic nucleus (STN), the external globus pallidus (GPe) with its prototypical and arkypallidal neuron populations [4, 5], and the substantia nigra. These circuits interact through what are often called the direct and indirect pathways, resulting in dynamic neural outputs that contribute to flexible motor control [6, 7]. Parkinson’s disease (PD) is characterized by the loss of dopaminergic neurons in the substantia nigra, leading to motor symptoms including tremor, rigidity, and bradykinesia [8]. A hallmark of the disease is the emergence of abnormal beta oscillations (13-30 Hz) within basal ganglia circuits [9], which correlate strongly with motor symptom severity and are suppressed by effective deep brain stimulation [10–13]. Recent insights reveal that pathological beta manifests as intermittent “beta bursts” that correlate more strongly with symptoms than do average beta power [14, 15].

Two competing hypotheses have emerged regarding mechanisms of beta generation in the basal ganglia. The first proposes that STN-GPe interactions form a critical pacemaker unit [16], supported by evidence that subthalamic activity correlates strongly with downstream basal ganglia activity in parkinsonian conditions [17, 18]. This excitatory-inhibitory network architecture is well-established to support oscillatory dynamics [19]. Foundational computational models demonstrate how subthalamopallidal networks may generate parkinsonian oscillations [20–25], with frequency related to slow potassium-dependent adaptation currents in the STN [26]. However, recent evidence has challenged the traditional STN-centric view, with opto-inhibition studies showing that STN suppression has minimal influence on GPe beta power [27]. As an alternative, experimental evidence from rodent studies suggests that pallidostriatal pathways may be the primary driver of beta oscillations [27–34], with computational models showing that these connections promote beta oscillations in dopamine-depleted networks [35]. Other studies have also implicated sites outside of the STN and GPe, namely cortical [36] and striatal [37, 38] circuits, in beta generation.

Most puzzling are contradictory results from recent modeling work. One computational study [39] demonstrated that dopamine depletion leads to pathological beta synchronization with STN activity playing an essential role for beta generation. Another from the same time frame [40] found that pallidostriatal circuit changes can trigger beta synchrony with STN actually weakening rather than promoting oscillations. This contradiction is particularly perplexing given that both studies employed similar circuit architectures and addressed the same fundamental question.

This work addresses these contradictory findings by systematically investigating how integrate-and-fire model selection influences simulated basal ganglia network dynamics. We demonstrate that differences in neuronal model implementation can fundamentally alter the predicted role of STN in beta generation, potentially explaining the conflicting results in recent literature. Our findings suggest that the mathematical structure underlying spike generation, rather than specific biophysical details, determines phase relationships and oscillation amplitudes within basal ganglia circuits.

The remainder of this paper is organized as follows. Section 2 introduces a simplified rate model to demonstrate how the phase relationship between STN and GPe prototypical neuron populations directly influences beta oscillation amplitude. Section 3 analyzes single neuron dynamics under periodic inhibition, using *time-to-spike functions* to prove that LIF and QIF neurons exhibit fundamentally different phase preferences: anti-phase and in-phase, respectively. Section 4 presents comprehensive spiking network simulations incorporating both pallidostriatal and subthalamopallidal circuits, demonstrating that the single-neuron phase preferences scale to population-level dynamics and determine whether STN enhances or suppresses beta oscillations. Section 5 discusses implications of our findings and directions for future research. Mathematical proofs supporting the analytical results are provided in the Appendix.

## Models and methods

All code for the models and simulations presented in this paper along with associated documentation is openly available in a GitHub repository at the following site: https://github.com/kanaptse/BG-Oscillation-Synergy-Suppression.

### Population firing rate model

We use a population firing rate model to provide a simplified setting in which to demonstrate the effect of phase relationships on beta oscillations. Let *P* = {STN, Proto, FSI, D2} be the set containing all populations in the model, which are subthalamic nucleus neurons, prototypical neurons of the external segment of the globus pallidus or GPe, striatal fast spiking interneurons, and striatal spiny projection neurons with D2 dopamine receptors (the predominant source of striatal inputs to the GPe), respectively. The rate dynamics for population *p* ∈ *P* obeys the equation

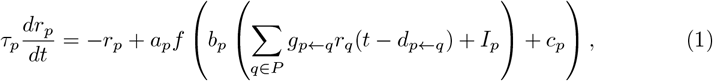

where *f* is the sigmoid function *f* (*x*) = 1*/*(1 + *e*^−*x*^). The population-specific parameters that we use in equation (1) are provided in Table 1, while connection strengths are detailed in Table 2. Connection strengths not explicitly listed in the table are set to zero. All parameters have been calibrated to match the physiological firing rates of each population: STN [12–20] Hz [41], Proto [40–60] Hz [5], FSI [10–20] Hz [42, 43], and D2 [0.5–2.5] Hz [44]. Among all possible transmission delays, we specifically vary only *d*_STN←Proto_ to control the phase relationship between STN and Proto populations, the impact of which is the primary focus of this preliminary part of our study. All other delays are set to zero. It is important to note that this manipulation does not represent any physiological mechanism; rather, it serves as a computational tool to systematically explore how different phase relationships affect network dynamics. In reality, phase relationships in the basal ganglia are determined by many factors. In the Results section, we will examine how intrinsic neuronal dynamics can modulate these phase relationships.

**Table 1.**
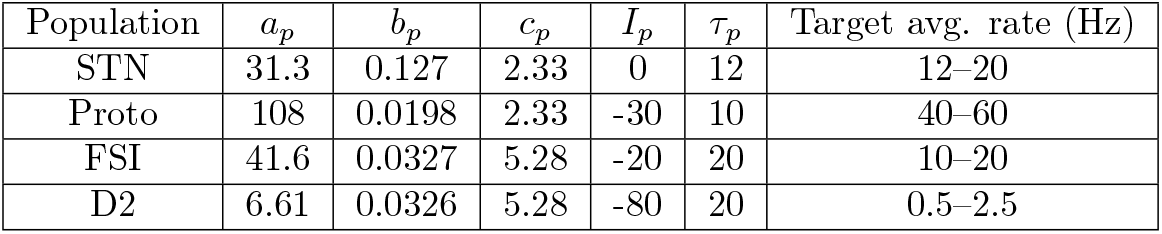
Parameter values for the firing rate model for the basal ganglia populations considered. The parameters *a*_*p*_, *b*_*p*_, *c*_*p*_, *I*_*p*_, and *τ*_*p*_ are shown for each neural population (STN, Proto, FSI, and D2), along with the target average firing rate ranges that these parameters were calibrated to achieve.

| Population | $a_p$ | $b_p$ | $c_p$ | $I_p$ | $\tau_p$ | Target avg. rate (Hz) |
| --- | --- | --- | --- | --- | --- | --- |
| STN | 31.3 | 0.127 | 2.33 | 0 | 12 | 12–20 |
| Proto | 108 | 0.0198 | 2.33 | -30 | 10 | 40–60 |
| FSI | 41.6 | 0.0327 | 5.28 | -20 | 20 | 10–20 |
| D2 | 6.61 | 0.0326 | 5.28 | -80 | 20 | 0.5–2.5 |

**Table 2.** Connection strengths between populations in the basal ganglia model. Each *g*_*p*←*q*_ represents the strength of the connection from population *q* to population *p*. These parameter values were selected to achieve the target firing rates specified in Table 1.

| Connection | Value |
| --- | --- |
| $g_{\text{STN} \leftarrow \text{Proto}}$ | 0.4 |
| $g_{\text{Proto} \leftarrow \text{D2}}$ | 50 |
| $g_{\text{Proto} \leftarrow \text{STN}}$ | 4 |
| $g_{\text{FSI} \leftarrow \text{Proto}}$ | 4 |
| $g_{\text{D2} \leftarrow \text{FSI}}$ | 10 |

### Phase difference analysis

Phase differences between Proto and STN populations in this paper were calculated using Fast Fourier Transform (`numpy.fft.fft`) analysis. The firing rate signals were demeaned and transformed to the frequency domain. The dominant frequency was identified as the frequency with maximum power in the Proto signal. Phase differences were calculated as the phase of STN minus the phase of Proto at the dominant frequency and expressed in degrees, wrapped to [-180°, 180°]. For each population, the phase at the dominant frequency (frequency with maximum power in the FFT power spectrum) was extracted from the complex FFT coefficient of the demeaned firing rate.

### Single neuron models

In examining the locking of single neurons to periodic inhibition, we use

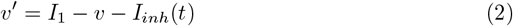

for leaky integrate-and-fire (LIF) neurons and

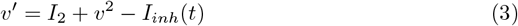

for quadratic integrate-and-fire (QIF) neurons. The inhibition function is defined as *I*_*inh*_(*t*) = *a* ·*H*(sin(2*πt/T*)), where *a* represents the inhibition amplitude, *H* denotes the Heaviside step function, and *T* is the inhibition period. Additionally, we implement boundary conditions such that *v*(0) = *v*_*reset*_ and *v*(*t*^+^) = *v*_*reset*_ when *v*(*t*^−^) = *v*_*th*_, where *v*_*reset*_ indicates the reset voltage and *v*_*th*_ represents the firing threshold. In this section, we set *v*_*reset*_ = −0.2 and *v*_*th*_ = 1.

### Time-to-spike function and map

We define a map, which we call the time-to-spike map [45], that we will use to analyze phase locking of a single neuron to a periodic signal. Suppose that square pulses of inhibition are delivered with period *T*, with inhibition on and off for equal times *T/*2 within each cycle. Let 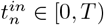 represent the time difference between the *n*-th firing event and the center of the subsequent inhibition period. We define the inhibition function by 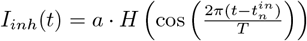, such that *t* = *t*_*in*_ represents the mid-point of the inhibition-on phase, as desired. The time-to-spike 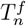 is determined by calculating the duration required for the dynamical system *v*^*′*^ = *f* (*v*) − *I*_*inh*_(*t*) to evolve from *v*_*reset*_ to *v*_*th*_. We express this relationship as 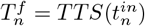, where *TTS* denotes the time-to-spike function.

To analyze the evolution of phase, we derive a map for 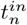. By definition, 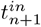 represents the time difference between the (*n* + 1)-st firing event and the center of the subsequent inhibition-on period. The mathematical formulation of this map takes the following form, which is illustrated schematically in Figure 1:

**Fig 1.**
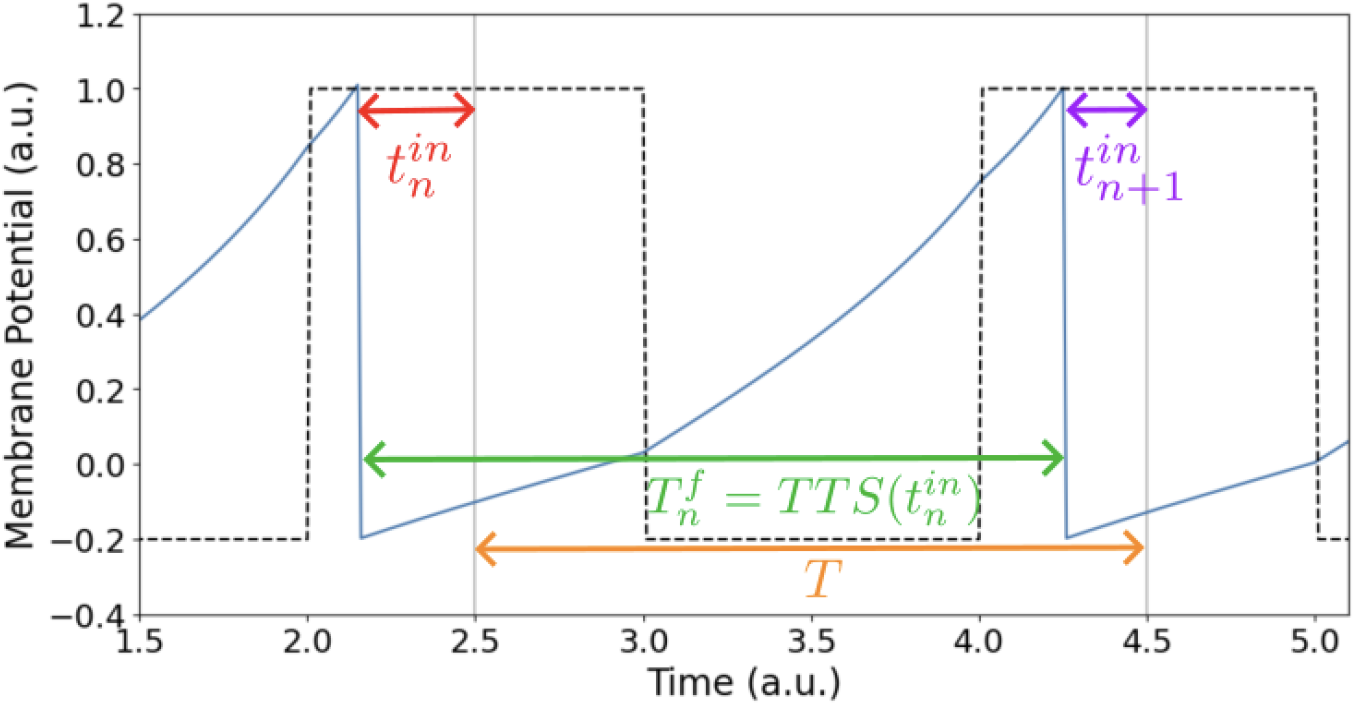
Schematic illustration of the time-to-spike (TTS) mapping. The diagram shows the relationship between consecutive firing events and inhibitory periods. The time difference between the *n*-th firing event and the center of the subsequent inhibition-on period, 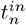, evolves to 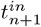 through the TTS function, defined to equal 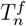, which is the time between the *n*-th and *n* + 1-st firing events. *T* is the period of inhibition.

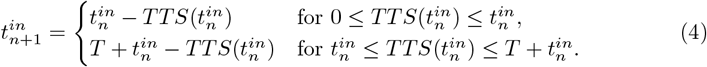

Note that this formulation is easily generalized to cases where there are multiple periods of inhibition between firing events, but we do not pursue that here.

### Multi-population basal ganglia model

We assembled collections of spiking neurons into a multi-population basal ganglia model, which we simulated. For these simulations, we use LIF or QIF equations to represent individual neurons. We specified these equations slightly differently than in the single-neuron case, to accommodate synaptic interactions, as follows.

#### Leaky integrate-and-fire (LIF) model

The membrane potential *V* of the LIF neuron evolves according to:

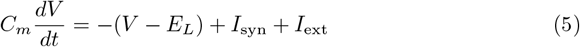

where *C*_*m*_ is the membrane capacitance, *E*_*L*_ is the leakage potential, *I*_syn_ is the synaptic current, and *I*_ext_ is the external current. When *V* reaches the threshold potential *V*_*th*_, the neuron fires a spike, and *V* is reset to *V*_*reset*_.

#### Quadratic integrate-and-fire (QIF) model

For the STN population in the alternative configuration, we used the QIF model:

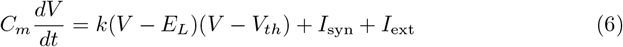

where *k* is a scaling factor. Note that in contrast to the LIF model, we use *V*_*th*_ here as a non-threshold parameter (see Table 3). As for threshold, when *V* reaches 0 mV, the neuron fires a spike, and *V* is reset to *V*_*reset*_.

**Table 3.** Population Parameters.

| Population | Model | Neurons | $C_m$ (pF) | $V_{th}$ (mV) | $E_L$ (mV) | $V_{reset}$ (mV) | $I_{ext}$ (pA) |
| --- | --- | --- | --- | --- | --- | --- | --- |
| D2 | LIF | 6000 | 4.9 | -50.0 | -76.8 | -76.8 | 24.8 |
| Proto | LIF | 780 | 12.9 | -54.8 | -65.0 | -65.0 | 16.0 |
| FSI | LIF | 420 | 3.1 | -52.4 | -78.2 | -78.2 | 25.5 |
| STN (LIF) | LIF | 408 | 5.1 | -50.8 | -59.0 | -59.0 | 9.1 |
| STN (QIF) | QIF | 408 | 20.0 | -54.0 | -70.0 | -70.0 | 66.0 |

To investigate how non-identical neurons affect population dynamics, we introduced heterogeneous external input currents for a parameter sweep analysis (Fig 13). Each neuron *i* received an external input current drawn from a Gaussian distribution:

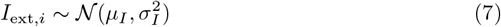

where *µ*_*I*_ is the mean input current for the population and *σ*_*I*_ = 0.1*µ*_*I*_ for LIF neurons and *σ*_*I*_ = 0.01*µ*_*I*_ for QIF neurons. All other simulations used homogeneous populations where each neuron within a population received identical external input currents. This heterogeneity allowed us to examine how neuronal variability within populations influences the phase coupling between STN and Proto and the resulting beta oscillation dynamics.

Synaptic interactions were modeled using exponential synapses with a time constant *τ* :

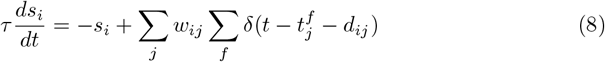

where *s*_*i*_ is the synaptic gating variable of the neuron *i, w*_*ij*_ is the connection weight from neuron *j* to neuron *i*, 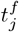 is the firing time of the presynaptic neuron *j*, and *d*_*ij*_ is the synaptic delay in the connection from neuron *j* to neuron *i*. The synaptic current is given by *I*_syn,*i*_ = *gs*_*i*_, where *g* is the maximal synaptic conductance.

Table 3 shows the parameters for each neuronal population, and Table 4 shows the connection parameters between populations. Parameters are adapted from [39, 40]. Negative strength values indicate inhibitory connections, while positive values indicate excitatory connections.

**Table 4.** Connection Parameters.

| Source | Target | Probability | Delay (ms) | $g$ (nS) | $\tau$ (ms) |
| --- | --- | --- | --- | --- | --- |
| D2 | Proto | 0.0833 | 7.0 | -0.460 | 5.5 |
| FSI | D2 | 0.0524 | 1.7 | -1.216 | 10.0 |
| Proto | FSI | 0.0128 | 7.0 | -1.170 | 5.5 |
| Proto | STN | 0.0385 | 1.0 | -0.096 | 8.0 |
| STN | Proto | 0.0735 | 2.0 | 0.061 | 10.0 |

We analyzed network dynamics using the following measures:

#### Population firing rate

For each population, the instantaneous firing rate was calculated by concatenating all spike times within the population and binning them with a time window of 2 ms. The firing rate was computed as the number of spikes per bin divided by the bin width and the number of neurons in the population, expressed in Hz.

#### Power spectral density

Power spectra were computed using Welch’s method (`scipy.signal.welch`) with a segment length of 256 samples and default overlap. The sampling frequency was determined from the bin width (500 Hz for 2 ms bins).

#### Beta power quantification

Beta power was defined as the average power spectral density within the beta frequency band (12.5-30 Hz). For each simulation, beta power was calculated from the population firing rate time series using `scipy.signal.welch` with nperseg=256. An initial transient period of 1000 ms was excluded to ensure steady-state dynamics.

#### Spectrograms

Time-frequency analysis was performed using `scipy.signal.spectrogram` with a segment length of 256 samples and overlap of 128 samples. The firing rate signal was demeaned by subtracting the mean before spectrogram computation to remove the DC component.

## Results

### Effect of phase relationship on oscillation amplitude in a rate model

To investigate how STN neuron dynamics affects beta oscillations in the broader basal ganglia network, we developed simplified rate models and spiking networks adapted from more complex, previously published computational models [39, 40]; see the previous section for details. Our network architecture (Fig 2) incorporates both the pallidostriatal and subthalamopallidal circuits, including four key neuronal populations: the excitatory subthalamic nucleus (STN) and the inhibitory external globus pallidus prototypical neurons (Proto), striatal D2-type spiny projection neurons (D2), and striatal fast-spiking interneurons (FSI). We focus on the subthalamopallidal (STN↔Proto) and pallidostriatal (D2→Proto→FSI→D2) loops, excluding arkypallidal neurons. This simplification allows us to isolate the core contradiction between recent studies: [39] showed that STN is necessary for beta oscillations generated by these coupled loops, while [40] found that STN suppresses them. Although arkypallidal projections can contribute to beta dynamics, they are not necessary for the fundamental disagreement we aim to resolve regarding how STN-Proto interactions modulate oscillations of pallidostriatal origin.

**Fig 2.**
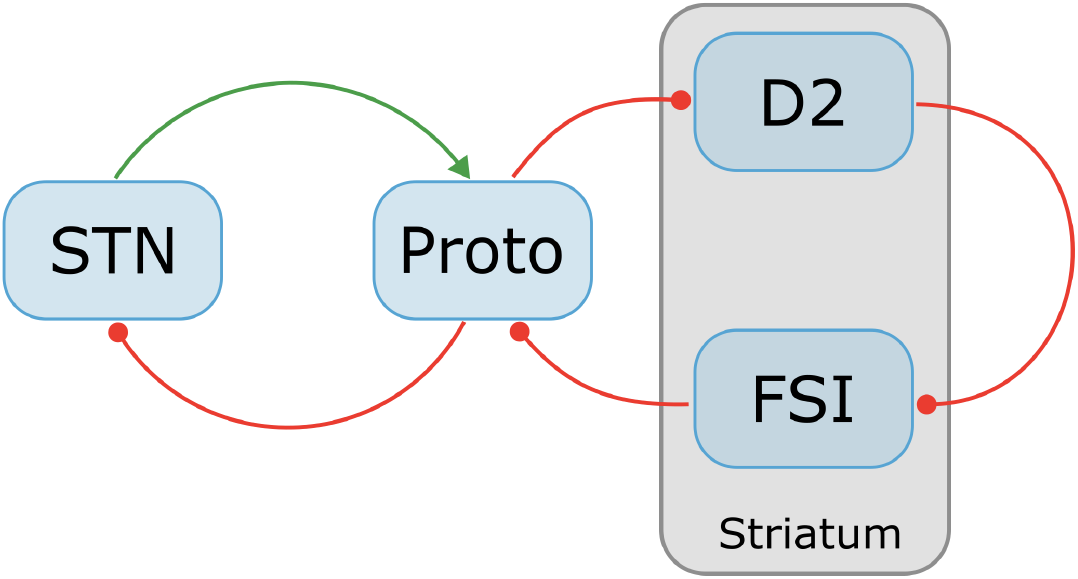
Schematic diagram of the basal ganglia circuit model showing four neuronal populations: subthalamic nucleus (STN), external globus pallidus prototypical neurons (Proto), striatal D2-type spiny projection neurons (D2), and striatal fast-spiking interneurons (FSI). Red arrows indicate inhibitory connections and the green arrow indicates an excitatory connection.

Recurrent connections within the inhibitory populations were omitted for simplicity, while experimental evidence suggests an abscence of recurrent excitatory connections within the STN population. We initialize the model such that the pallidostriatal loop (D2→Proto→FSI→D2) generates baseline beta oscillations, while the subthalamopallidal circuit (STN↔Proto) provides modulatory input. In this section, we will show how the oscillation amplitude in this coupled network depends on the phase relationship between STN and Proto firing rates.

To investigate how phase relationships between STN and Proto populations affect oscillation amplitude, we first examine some specific example cases in a simplified firing rate model. We manipulate the delay from Proto to STN to control their phase relationship. In Figure 3, the Proto-FSI-D2 circuit is configured to generate a stable periodic oscillation. The STN to Proto connection is activated at t = 975 ms to examine how STN input modulates the ongoing oscillation. The results of 3 cases are shown: without STN, with STN but no delay, and with both STN and delay. Without delay, the two populations’ firing rates show an anti-phase relationship (Fig 3B). As we add sufficient delay the two populations switch to an in-phase relationship (Fig 3C, 16 ms delay). Notably, the introduction of STN with no delay (Fig 3A, green curve) decreases the Proto oscillation amplitude compared to the baseline condition (blue curve), while the addition of delay (red curve) substantially increases the amplitude. These amplitude variations can be understood through the diagram in Figure 3D, which illustrates the trajectory of oscillations in the state space. For the no-delay case, the STN excitation arrives during the Proto activity trough, reducing its depth and dampening the overall oscillatory behavior. Conversely, in the with-delay condition, the excitation aligns with the rising phase of Proto activity, reinforcing and amplifying the oscillations.

**Fig 3.**
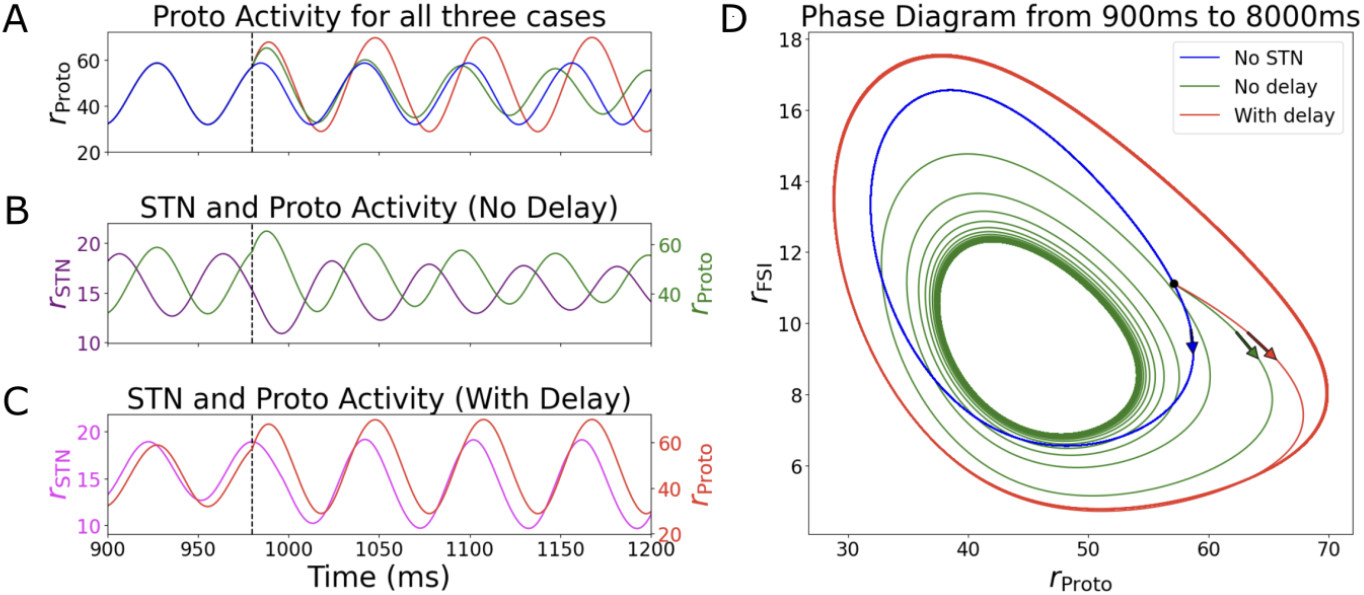
Rate model simulations demonstrating how STN-Proto phase relationships modulate oscillation amplitude. The STN to Proto connection is activated at *t* = 975 ms. (A) Proto population activity under three conditions: baseline without STN (blue), STN input with no delay (green), and STN input with 16 ms delay (red; in this case, the input to STN is based on Proto activity 16 ms earlier). (B) Anti-phase relationship of STN and Proto firing rates in the no-delay condition. (C) In-phase STN-Proto phase relationship with 16 ms delay. (D) Trajectories in state space (900–8000 ms) illustrating the mechanistic basis for the amplitude effects: anti-phase excitation arrives during Proto activity troughs creating smaller orbits (green), while in-phase excitation aligns with rising Proto activity producing larger orbits (red). Arrows highlight divergence of trajectory paths following STN activation.

To systematically investigate how the phase relationship between STN and Proto populations modulates beta oscillation amplitude in Proto, we conducted a parameter exploration study. Figure 4 presents the results of this analysis. Figure 4A shows oscillation amplitude as a function of the drive to Proto (*I*_Proto_) and the STN to Proto connection strength (*g*_Proto←STN_) without any delay. Varying *I*_Proto_ for fixed *g*_Proto←STN_ below approximately 3.5 reveals two presumed Andronov-Hopf (AH) bifurcations, as evidenced by the emergence and subsequent disappearance of oscillations along horizontal sections of the heatmap. However, the vertical patterns highlight a key difference: for fixed *I*_Proto_ values in the range of −60 to −20, oscillation amplitude decreases monotonically as *g*_Proto←STN_ increases. Notably, increasing *g*_Proto←STN_ increases the excitatory drive to Proto, but unlike increasing *I*_Proto_, it does not cause an initial increase in oscillation amplitude before the decrease. This confirms that the decrease in amplitude with small *g*_Proto←STN_ is not caused by the occurrence of a second Hopf bifurcation.

**Fig 4.**
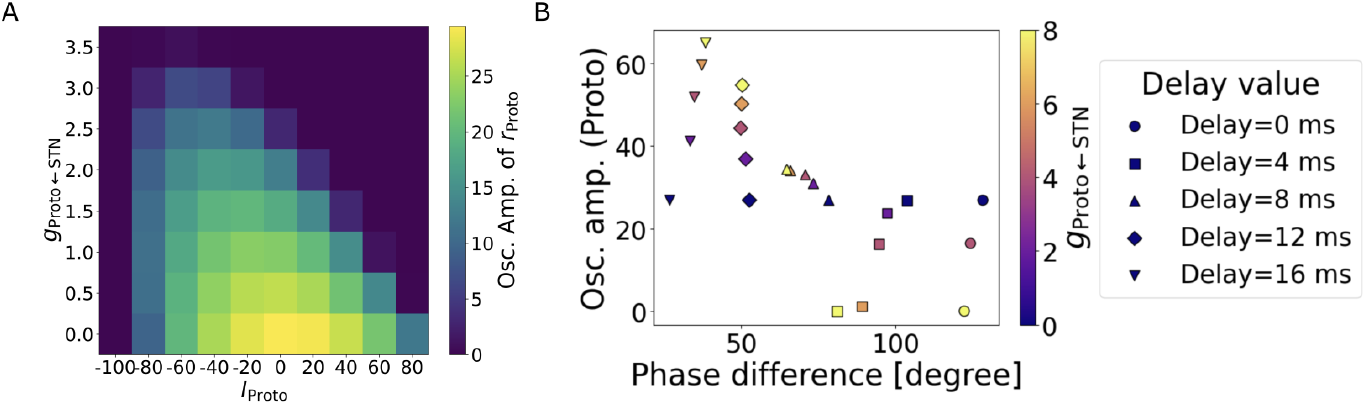
Parameter exploration of factors affecting beta oscillation amplitude in the STN-Proto circuit. (A) Heat map showing oscillation amplitude (color) as a function of STN to Proto connectivity strength (*g*_Proto←STN_) and drive to Proto (*I*_Proto_). The amplitude decreases with increasing connectivity strength across the full range of drive values, including regions between the two Hopf bifurcations, demonstrating that the amplitude reduction is not attributable to the second Hopf bifurcation. (B) Scatter plot depicting the relationship between phase difference and oscillation amplitude across different Proto to STN delay values (*d*_Proto←STN_). Each point represents a different connectivity strength (*g*_Proto←STN_, color-coded), with symbols indicating delay values (circles: 0 ms, squares: 4 ms, triangles: 8 ms, diamonds: 12 ms, inverted triangles: 16 ms). The plot reveals that shorter delays produce larger phase differences (anti-phase, 90+ degrees) with lower amplitudes, while longer delays generate smaller phase differences (in-phase, ~20-60 degrees) with higher amplitudes.

Figure 4B illustrates the relationship between phase difference and oscillation amplitude across different delay and connectivity conditions. The scatter plot reveals a clear inverse relationship: as phase differences decrease (indicating a shift toward in-phase dynamics), oscillation amplitudes increase substantially. At short delays (0-4 ms, circles and squares), the STN and Proto populations exhibit large phase differences of approximately 100-120 degrees, corresponding to anti-phase relationships that suppress oscillatory activity and result in low amplitudes (typically below 30). In this case, larger *g*_Proto←STN_ values lead to reduced oscillation amplitudes. Conversely, at longer delays (12-16 ms, diamonds and inverted triangles), phase differences decrease to 20-60 degrees, indicating in-phase relationships that enhance oscillatory activity and produce amplitudes reaching 50-60. Here, greater *g*_Proto←STN_ strengths yield enhanced oscillation amplitudes. The intermediate delay condition (8 ms, triangles) shows a transitional behavior with moderate phase differences around 60-80 degrees and intermediate amplitudes, with less dependence on *g*_Proto←STN_.

Together, these results provide a comprehensive view of how both connectivity strength and relative timing of STN and Proto oscillations, dictated by the delay in input from STN to Proto, contribute to the generation and modulation of beta oscillations in the basal ganglia circuit.

### Single neuron dynamics and phase relationships under periodic inhibition

Having established the influence of phase relationship on oscillation amplitude, we next asked what could cause different phase relationships to materialize. We hypothesized that within a fixed circuit structure, the choice of neuronal model used would determine the phase differences that emerge. Our goal in this section is to understand how different models could yield distinct phase relationships between coupled excitatory and inhibitory populations. We define *t*^*in*^ to be the difference in times of firing rate peaks of the two populations, as depicted by the first panel of Figure 5. Initially, we consider a highly simplified scenario in which we replace the inhibitory population and its firing rate with a periodic train of square pulses of inhibition, with inhibition on for half of each period and off for the other half, and the excitatory population and its firing rate with the voltage and spike times of a single excitatory neuron. In this setting, *t*^*in*^ is defined to be the time difference between the spike of the excitatory neuron and the mid-point of the time that its inhibitory input is on within each cycle, as depicted by the second panel of Figure 5. We will perform some computations and analysis of phase locking in this simplified setting, and later we will show with computational results that networks of spiking neurons exhibit similar properties to the simplified system.

**Fig 5.**
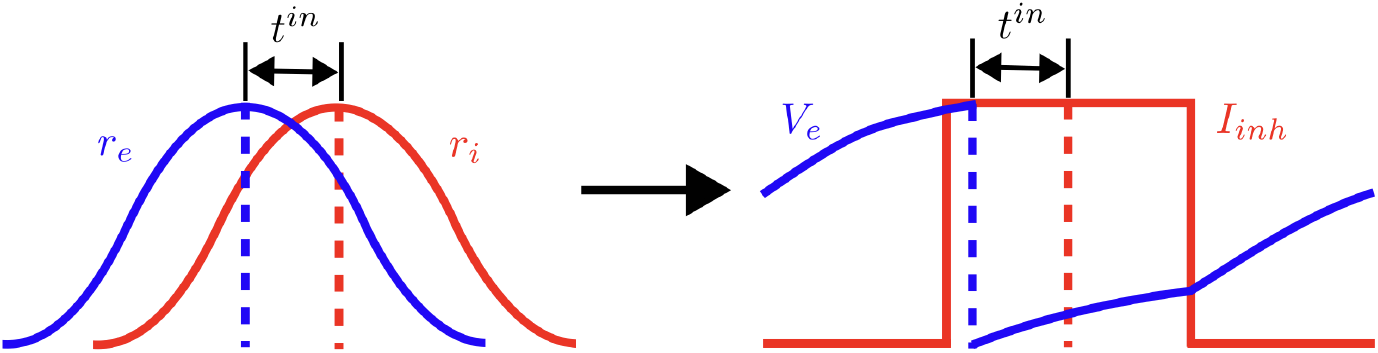
Schematic illustration of a simplified modeling approach for analyzing phase relationships between excitatory and inhibitory populations. The figure shows the transition from the original definition of *t*_*in*_ as the time difference between the peaks of excitatory (E) and inhibitory (I) population firing rates (left) to the simplified single neuron model where *t*_*in*_ is redefined as the temporal offset between the excitatory neuron’s spike times, where its voltage *V*_*e*_ reaches a maximum and is reset, and the mid-point of each pulse of square-wave inhibition that the excitatory neuron receives (right). This simplification preserves the essential phase locking effect while allowing for an analytical treatment of the underlying mechanisms.

#### Examples of phase relationships

We next present examples illustrating different phase relationships between neuronal firing and inhibitory input in QIF and LIF neurons (see Methods section). We demonstrate a fundamental difference in phase preferences: QIF neurons tend to fire during inhibition periods (in-phase), while LIF neurons preferentially fire when inhibition is absent (anti-phase).

Figure 6 provides a visual representation of these contrasting behaviors. In the QIF model (panels A and B), the membrane potential (green curve) evolves so that, after a few transient cycles, it reaches the firing threshold (*v* = 1) during periods of inhibition (black dashed). Following each threshold crossing, the potential is reset to *v*_*reset*_ = − 0.2. Conversely, in the LIF model (panels C and D), threshold crossings occur during periods when inhibition is off. The phase plane representations in panels B and D reveal the underlying dynamical structure responsible for these different behaviors, which will be analyzed formally in subsequent sections. We have chosen parameters so that the neurons are phase-locked to the periodic inhibition, and they converge to steady-state periodic behavior over time.

**Fig 6.**
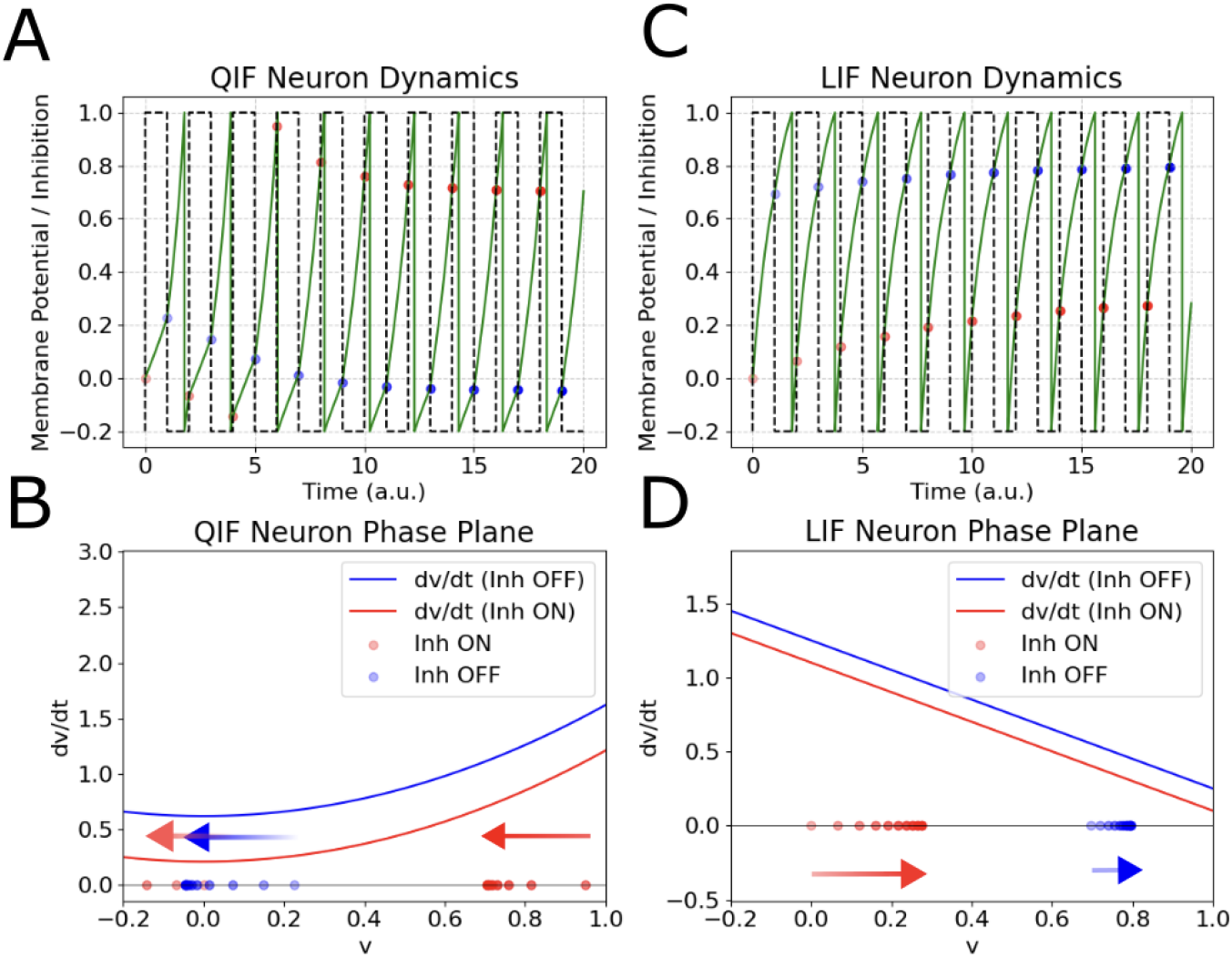
Comparison of QIF and LIF neuron dynamics under square-wave inhibitory forcing. (A) Membrane potential dynamics of a QIF neuron. Red dots indicate moments when inhibition turns ON, blue dots when inhibition turns OFF. (B) Phase plane representation of the QIF neuron, with membrane potential (*v*) on the x-axis and *dv/dt* on the y-axis. Dots correspond to the same time points shown in panel A, with matching transparency indicating simultaneous moments across the two panels. Lighter dots correspond to earlier times, and the arrows indicate the chronological order of the blue and red dots, respectively. (C) and (D) are similar figures for an LIF neuron. Note how QIF neurons preferentially fire during inhibition (in-phase relationship), while LIF neurons fire predominantly when inhibition is absent (anti-phase relationship).

In all four panels of Figure 6, red and blue dots indicate the neuron’s voltage when inhibition turns on and off, respectively. The transparency of the dots, mostly visible in panels B and D, reflects their temporal occurrence, with higher transparency (paler) dots indicating earlier time points in the simulation. For QIF neurons, the voltage is typically low (near 0) when inhibition is off and high when inhibition is on. LIF neurons exhibit the opposite pattern. The arrows in panels B and D illustrate the convergence pattern of these voltage states toward their steady-state values. These observations provide mechanistic insight into why the two neuron types exhibit different phase relationships. QIF neurons have a saddle-node bifurcation giving rise to two low-voltage equilibria as their inhibition level increases. Whether inhibition is on or off, *dv/dt* is small near the “ghost” of these equilibria, but this effect is especially strong with inhibition present. Thus, if the neuron fires with inhibition off on one cycle, then its voltage stays close to the reset value throughout the inhibition-on period and then does not have enough time to evolve up to the firing threshold when inhibition is off, resulting in an inhibition-on spike on the next cycle. From another perspective (Fig 6 B), the voltage levels at which inhibition turns on and off show phase precession until they stabilize with low (high) voltages when inhibition turns off (on). In contrast, in LIF neurons, the linear form of the *v*-nullcline allows voltage to rise significantly during inhibition yet slows it near threshold, thereby suppressing firing. When inhibition is removed, the increase in *dv/dt* allows a faster increase in *v* that facilitates threshold crossing. In this case, voltages when inhibition turns on and off show phase advances and stabilize with high (low) voltages when inhibition turns off (on).

We will next analyze these effects using the *time-to-spike* (TTS) function to provide a mathematical proof demonstrating why in-phase solutions cannot be stable for LIF neurons and confirming the existence of stable in-phase solutions for QIF neurons.

#### Time-to-spike analysis: fixed points and phase relationships

We defined a time-to-spike function (4) [45] that serves as a map from the time when a periodically driven neuron spikes on one input cycle, relative to a reference time defined based on the drive signal, to the corresponding relative time on the next input cycle (see Methods). Suppose that a fixed point 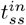 of the map (4) exists. An anti-phase steady state corresponds to 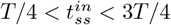, where neurons fire during the inhibition-off period, while an in-phase solution corresponds to 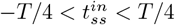, where neurons fire during inhibition.

We focus on phase-locked solution corresponding to the *k* = 1 case of equation (4). To solve for the fixed point 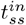, we substitute 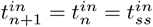 into the equation. Hence, we have 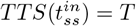. For the fixed point to be stable, we need the derivative on the right-hand side to be between − 1 and 1. Hence, we require 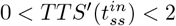.

Figure 7 illustrates the numerically computed TTS functions and their corresponding fixed points for both LIF and QIF neurons with a period of *T* = 2 ms, set by the period of the inhibition function. For LIF neurons (panels A-B), our numerics reveals a stable fixed point, which we will denote as 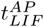, with a distance above *T/*4 = 1*/*2 from the inhibition center (green circle) corresponding to anti-phase firing during inhibition-off periods. Additionally, an unstable fixed point 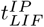 (red circle) exists at a distance less than *T/*4 from the inhibition center, corresponding to in-phase firing.

**Fig 7.**
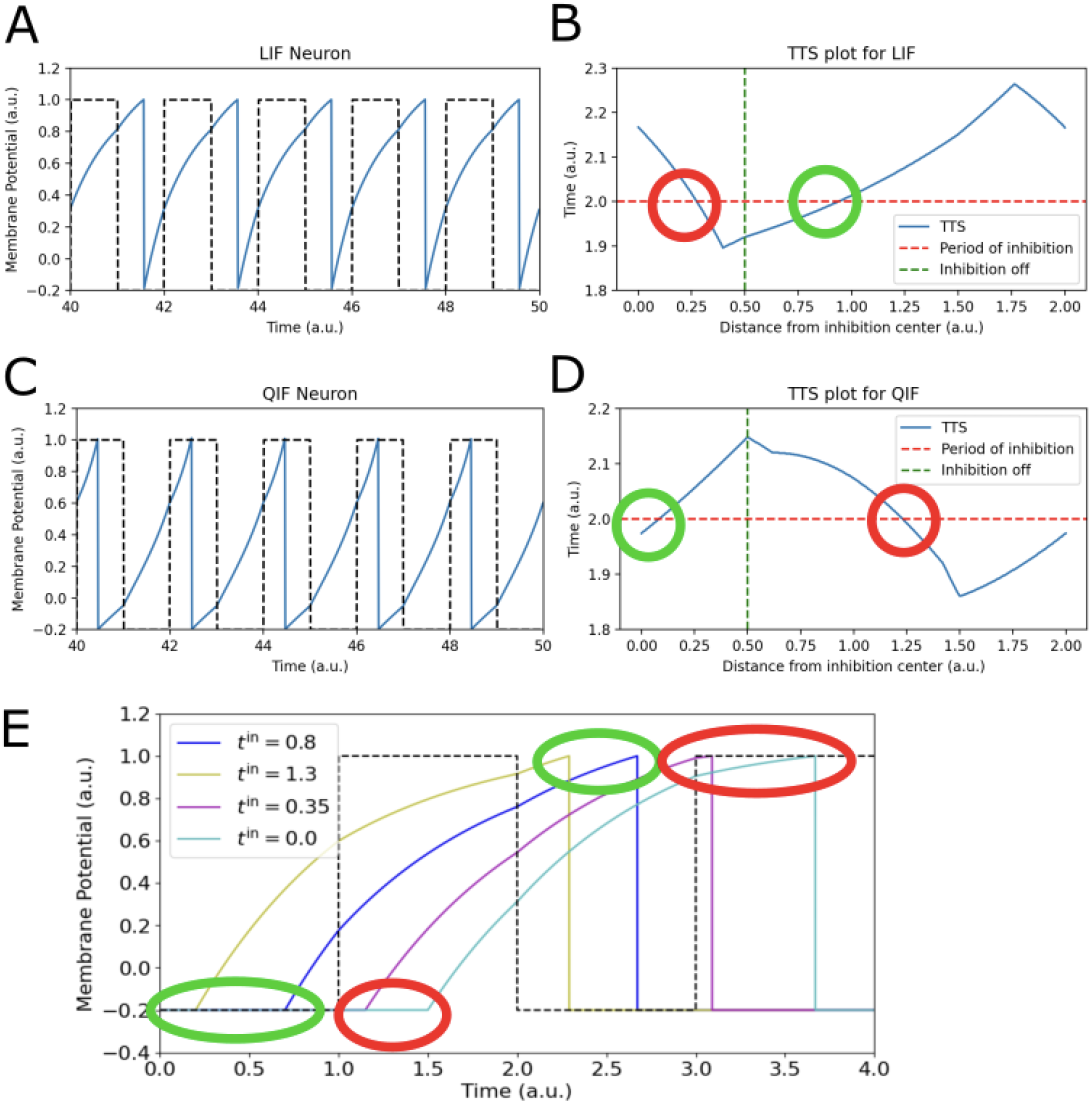
Numerics of Time-To-Spike (TTS) functions and fixed points for LIF and QIF neurons with a period of 2 ms. (A) Phase relationship in LIF neurons showing predominant firing during inhibition-off periods (anti-phase). (B) TTS function for LIF neurons displaying stable anti-phase 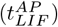 and unstable in-phase 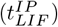 fixed points. (C) Phase relationship in QIF neurons showing predominant firing during inhibition-on periods (in-phase). (D) TTS function for QIF neurons with opposite fixed point stability. (E) Single-step TTS mapping for LIF neurons demonstrating regions of initial conditions that lead to contraction (green) and dilation (red).

Conversely, QIF neurons (panels C-D) exhibit the opposite behavior, with a stable fixed point, 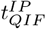, corresponding to in-phase firing and an unstable fixed point, 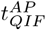, corresponding to anti-phase firing. This fundamental difference in dynamics agrees with and provides mathematical support for the contrasting phase relationships observed in Figure 6.

The stability properties of these fixed points can be understood from the slope of the TTS function at each intersection with the *TTS*(*t*^*in*^) = *T* line. Panel E demonstrates a contraction between trajectories that occurs on an interval of initial conditions of the LIF model (green region), constraining the slope of the TTS function such that 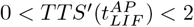 and hence leading to the stability of anti-phase solutions. Conversely, the red region represents a dilation giving rise to the slope 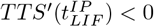 (red region), corresponding to the instability of in-phase solutions. These mathematical properties, formalized in the Appendix, provide a rigorous foundation for understanding the emergence of different phase relationships in these neuronal models.

To demonstrate the robustness of our findings across different parameter regimes, we conducted a numerical study of the steady-state phase relationships as a function of both inhibition amplitude and excitatory drive strength. Figure 8 presents the results for both LIF and QIF neurons under periodic inhibition. For each parameter combination, we numerically computed the time-to-spike function *TTS*(*t*^*in*^) by integrating the voltage equation across a grid of initial phase values. We then identified stable fixed points satisfying 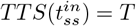 with 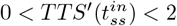 by detecting positive-slope crossings of the fixed point condition. Phase values for these stable fixed points were calculated as 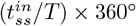, with color-coded regions displayed only where such stable fixed points exist.

**Fig 8.**
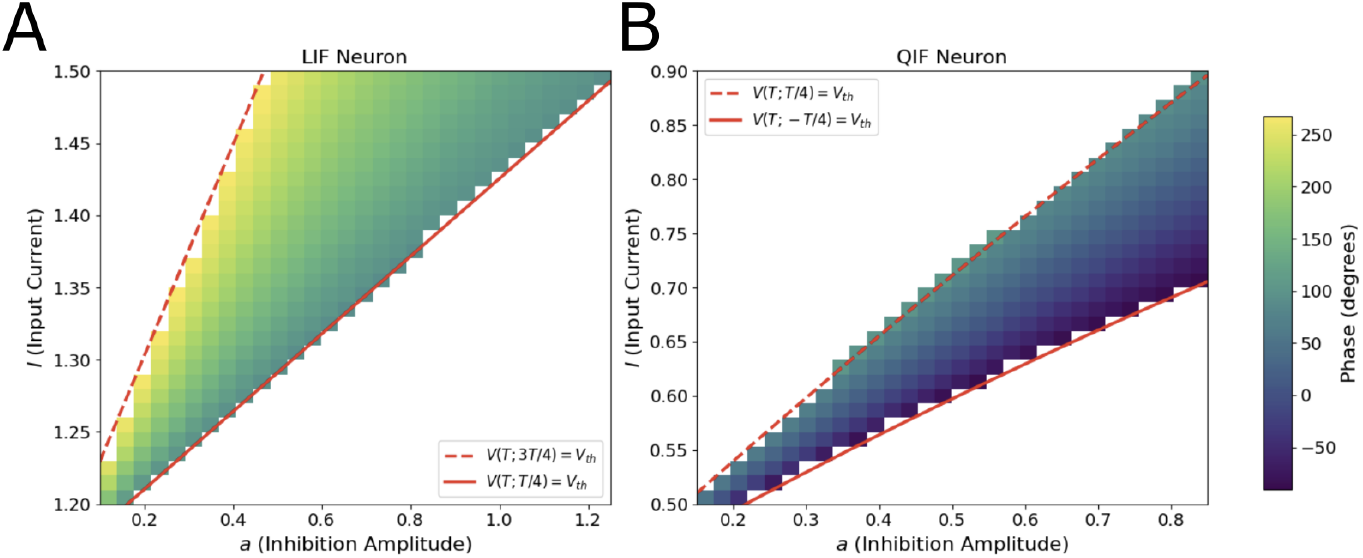
Steady-state phase relationships between STN and Proto populations as a function of excitatory drive *I* and inhibition amplitude *a*. (A) LIF neurons with *V*_reset_ = 0 exhibit phase relationships ranging from 90^◦^ (anti-phase, blue) to 270^◦^ (predominantly anti-phase, yellow). (B) QIF neurons display phase values from −90^◦^ to 130^◦^, showing stronger tendency toward in-phase relationships. Red lines show theoretical boundaries where phase-locking ceases to exist, derived from existence conditions (see Appendix). Between these boundaries, neurons maintain stable phase-locking to periodic inhibition, with the steady-state phase varying smoothly with parameters (indicated by color). The close agreement between analytical predictions (red lines) and numerically observed boundaries validates the time-to-spike framework. For LIF neurons, the boundaries are *V* (*T*; *T/*4) = *V*_th_ (solid) and *V* (*T*; 3*T/*4) = *V*_th_ (dashed). For QIF neurons, the boundaries are *V* (*T*; *T/*4) = *V*_th_ (dashed) and *V* (*T*; −*T/*4) = *V*_th_ (solid). Parameters: *T* = 2, *V*_th_ = 1.0, *V*_*r*_ = 0 (LIF), *V*_*r*_ = − 0.2 (QIF).

LIF neurons (Fig 8A) exhibit phase values predominantly ranging from 90^◦^ to 270^◦^, demonstrating their general preference for anti-phase relationships across most parameter combinations where stable solutions exist. In contrast, QIF neurons (Fig 8B) show a phase range from − 90^◦^ to 130^◦^, with most values in the middle part of this range, indicating a strong and consistent tendency toward in-phase relationships.

To understand these transitions analytically, we derived existence conditions for phase-locked solutions (see Appendix). These conditions predict sharp boundaries in parameter space: one boundary marks where phase-locking ceases to exist (when *V* (*T*; *T/*4) = *V*_th_), while the other marks the transition from anti-phase to in-phase relationships (when *V* (*T*; −*T/*4) = *V*_th_). The theoretical predictions (red lines in Figure 8) closely match the numerically observed phase transitions, confirming that our time-to-spike analysis accurately captures the mechanisms governing phase relationships.

This systematic exploration of parameter space demonstrates that the phase relationship preferences we characterized are not confined to specific parameter choices but represent fundamental properties of the respective model neuron types. The consistent differences in phase distributions—with LIF neurons showing broader anti-phase ranges and QIF neurons exhibiting tighter in-phase relationships—strengthen our conclusion that properties of intrinsic neuronal dynamics, rather than particular parameter tuning, determine phase relationships in oscillatory circuits.

#### Exponential integrate-and-fire neurons

Our investigation into how neuronal model choice affects beta oscillation dynamics was originally motivated by the computational study of [39], which demonstrated that dopamine depletion leads to pathological synchronization in the beta band, with their results suggesting that STN activity is essential for beta generation. Notably, their model implemented STN neurons using the exponential integrate-and-fire (EIF) framework rather than the QIF neurons we analyzed in previous sections. To verify that our conclusions regarding phase relationships and beta modulation hold for the original neuronal model used in their study, we examined EIF neuron dynamics under periodic inhibition.

The EIF model provides an intermediate level of biophysical realism between simple integrate-and-fire models and full conductance-based descriptions. It incorporates an exponential term that captures the rapid depolarization near spike threshold observed in real neurons:

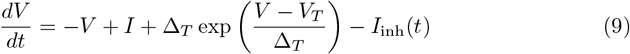

where Δ_*T*_ = 0.14 mV is the slope factor, *V*_*T*_ = 0.0 mV is the threshold potential, *I* = 0.14 pA is the external current, *I*_inh_(*t*) = −0.21 pA is the inhibitory current amplitude, *V*_th_ = 1.0 mV is the firing threshold, and *V*_reset_ = − 0.2 mV is the reset potential. The exponential term in (9) creates a sharp acceleration toward spike threshold that more closely approximates the sodium channel activation underlying action potential initiation than does the spiking behavior of the QIF.

Our simulations of EIF neurons under periodic inhibition display striking similarities to QIF neuron behavior, confirming that the conclusions drawn from [39] align with our theoretical framework. Figure 9 demonstrates that EIF neurons exhibit a clear preference for in-phase firing with periodic inhibition. The membrane potential dynamics (Fig 9A) show that spikes consistently occur during inhibition-on periods (red dots), with the neuron maintaining low membrane potentials during inhibition-off periods (blue dots), mirroring the pattern observed in QIF neurons (Fig 6A).

**Fig 9.**
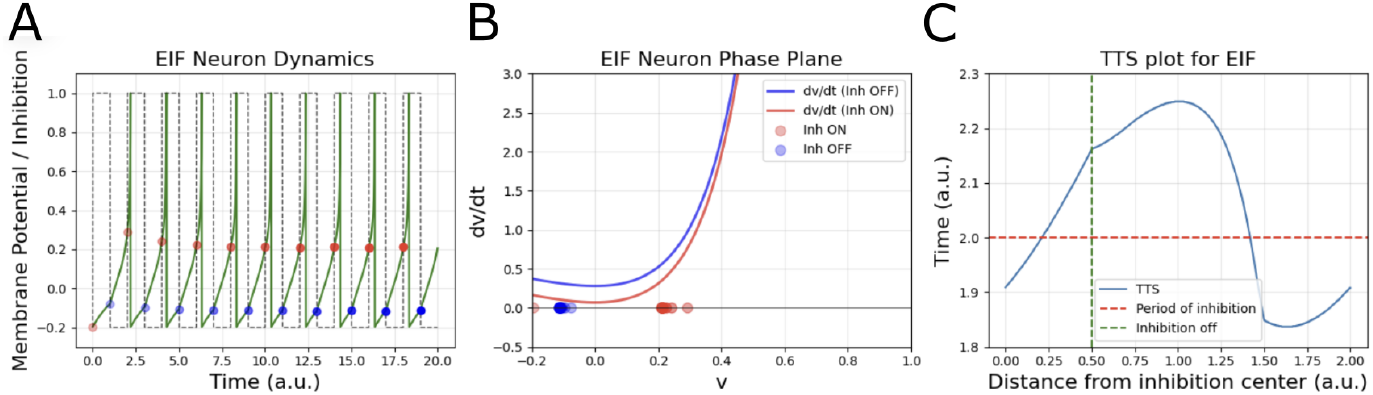
Exponential integrate-and-fire (EIF) neuron dynamics under periodic inhibition demonstrate in-phase firing preference similar to QIF neurons. (A) Membrane potential dynamics showing preferential firing during inhibition-on periods (red dots). (B) Phase plane representation showing rapid acceleration toward threshold during inhibitory periods (red) and low voltages while inhibition is off (blue). (C) Time-to-spike function with a stable (unstable) fixed point corresponding to an in-phase (anti-phase) relationship of neural firing relative to the inhibitory input, similar to patterns observed in Figure 7 for QIF neurons.

Phase plane analysis (Fig 9B) reveals the mechanistic basis for this in-phase preference. The *dV/dt* nullclines for inhibition-on (red) and inhibition-off (blue) conditions show that the exponential term in EIF neurons creates dynamics similar to those observed in QIF neurons (Fig 6B). During inhibition (red), the exponential acceleration still allows the neuron to reach threshold, while during inhibition-off periods (blue), the membrane potential tends to settle at lower values. The time-to-spike function for EIF neurons (Fig 9C) exhibits qualitative agreement with that for QIF neurons (Fig 7D), with a stable fixed point, 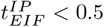, where the TTS curve intersects the inhibition period (red dashed line), corresponding to an in-phase solution.

These findings provide direct validation that the results of [39] are consistent with our theoretical predictions. Since EIF neurons exhibit in-phase coupling preferences similar to QIF neurons, STN populations modeled with EIF dynamics would establish in-phase relationships with Proto populations, thereby enhancing beta oscillations generated by the pallidostriatal circuit—exactly as observed in their study.

The similarity between EIF and QIF neuron dynamics in terms of phase relationships strengthens our conclusion that the mathematical structure underlying spike generation, rather than specific biophysical details, determines phase preferences under oscillatory conditions. Both models share the property of rapid, nonlinear approach to threshold, which enables firing during inhibitory periods when the neuron has sufficient momentum to overcome the additional hyperpolarizing drive.

This convergent behavior across QIF and EIF models suggests that STN neurons, which are known to exhibit rapid spike initiation kinetics, would likely demonstrate in-phase coupling with Proto populations during beta oscillations. This mechanistic insight provides additional support for the hypothesis that contradictory findings in the parkinsonian beta literature may stem from differences in neuronal model implementation rather than fundamental disagreements about circuit function, while simultaneously validating the theoretical framework underlying the findings of earlier simulations [39].

### Full neuronal network model

Up to this point, we have run simulations with firing rate models and with a spiking STN model coupled to a periodic inhibitory signal. To conclude our study, we next simulated a multi-population basal ganglia network model composed of spiking neurons with interconnections matching the known neuroanatomy (see Methods). We simulated four key populations: D2-type spiny projection neurons (D2), prototypical neurons in the external globus pallidus (Proto), fast-spiking interneurons (FSI), and subthalamic nucleus neurons (STN). The network topology follows the indirect pathway of the basal ganglia, which is known to be involved in beta oscillation generation.

#### Comparison between single neuron analysis and spiking network dynamics

To validate the biological relevance and broader applicability of our single-neuron analysis, we first implemented spiking network models consisting of interconnected STN and Proto populations, without the striatal elements from Figure 2. Figure 10 presents simulation results from these networks under periodic inhibitory drive to the Proto population, mimicking optogenetic stimulation protocols used in experimental settings [27].

**Fig 10.**
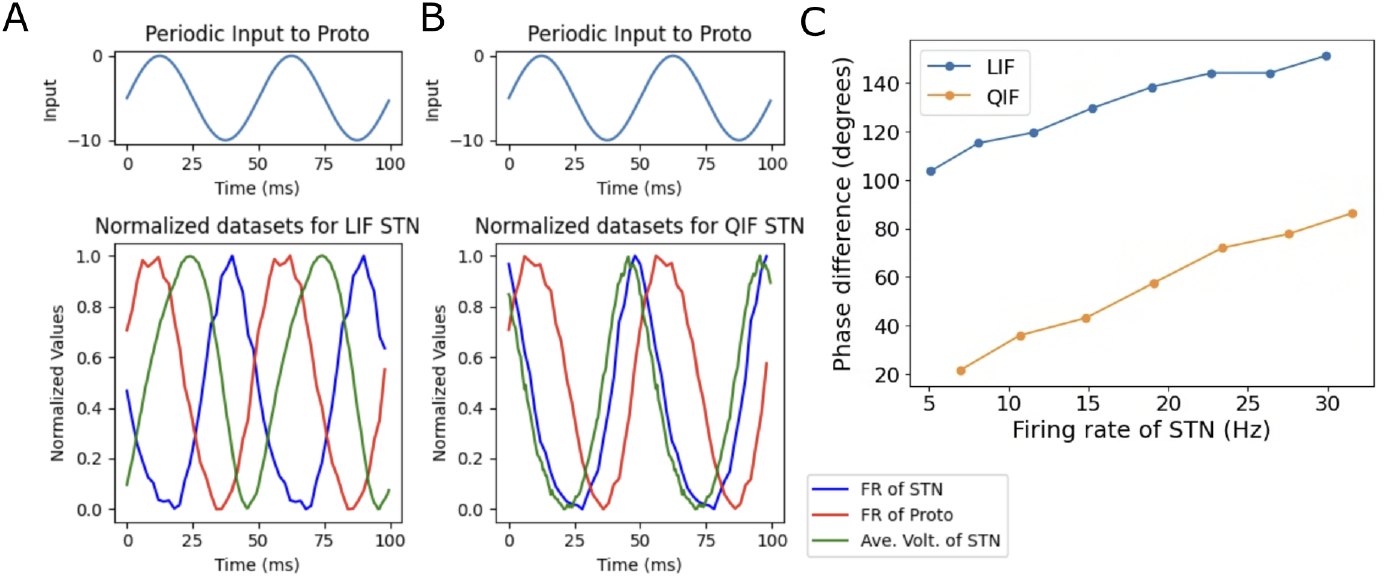
Comparative dynamics of STN-Proto circuits using different neuronal models. (A) Periodic inhibitory input to Proto neurons (top) and in-phase synchronization between QIF STN and Proto firing rates (bottom), consistent with single neuron studies. (B) Periodic inhibitory input to Proto neurons (top) and anti-phase relationship between LIF STN and Proto firing rates (bottom), with STN membrane potential continuing to rise up toward threshold even while Proto firing ramps up. In both (A) and (B), the Proto population consists of LIF model units. (C) Phase difference as a function of firing rate for LIF (blue) and QIF (orange) STN neurons, relative to Proto firing. LIF neurons consistently exhibit larger phase differences (anti-phase preferences), while QIF neurons show smaller phase differences (in-phase tendencies) across the firing rate spectrum.

In both network configurations, Proto neurons were modeled as LIF neurons, while STN neurons were implemented as either QIF (Fig 10A) or LIF (Fig 10B) neurons. This allowed us to isolate the effects of excitatory neuron dynamics on network-level phase relationships. The simulations incorporated full connectivity between populations with synaptic delays and conductance-based synapses.

The network-level simulations demonstrate striking concordance with our single-neuron predictions. In the LIF-STN network (Fig 10A), both the STN population firing rate (blue) and average membrane potential (green) exhibit clear anti-phase relationships with the Proto population activity (red), confirming our analytical prediction that LIF neurons tend toward anti-phase locking with periodic inhibition. Notably, the STN membrane potential dynamics (green curve) are consistent with the single-neuron phase plane analysis shown in Figure 6: membrane potentials are low during inhibition-on periods and elevated during inhibition-off periods, matching the voltage state patterns observed in our theoretical analysis.

Conversely, the QIF-STN network (Fig 10B) displays a pronounced in-phase relationship between STN and Proto population activities. Notably, the STN membrane potential rises substantially during periods of Proto inactivity (corresponding to inhibition-off periods in our single-neuron framework), such that STN is able to fire after Proto activity has risen from its trough, demonstrating the mechanistic basis for in-phase preferences in QIF neurons.

To further validate our theoretical predictions beyond the phase-locked regime, we examined how phase relationships vary as a function of population firing rate under periodic inhibition (see Methods). Figure 10C presents this analysis for both LIF and QIF neurons across a range of firing frequencies. The input current ranges for both model classes were selected to generate firing rates spanning from just above 0 Hz to approximately 30 Hz.

These results demonstrate that the phase preferences we identified persist across different firing rate regimes, even when neurons are not strictly phase-locked to the inhibitory input. LIF neurons (blue curve) consistently exhibit larger phase differences relative to the inhibitory input, confirming their anti-phase preference across the entire firing rate spectrum examined. Conversely, QIF neurons (orange curve) maintain smaller phase differences overall but exhibit a gradual increase in phase difference as firing rate increases. This trend can be explained by the increased input drive: as the excitatory drive to STN neurons is elevated, they eventually begin to fire more readily when inhibition level drops rather than maintaining their preferred in-phase timing.

This analysis extends our findings beyond the idealized phase-locked scenarios and demonstrates that the fundamental phase preferences determined by intrinsic neuronal dynamics remain consistent across physiologically relevant firing rate ranges. The persistence of these phase relationships across different firing frequencies strengthens our conclusion that neuronal model choice critically determines phase dynamics in oscillatory neural circuits.

These simulation results, including both the network dynamics and firing rate analysis, confirm that the phase relationships established in our single-neuron analysis extend robustly to the population level, even in the presence of network-level complexities. The consistency between single-neuron predictions and network behavior underscores the critical role of intrinsic neuronal dynamics in determining population-level synchronization patterns in the STN-Proto circuit.

Moreover, these findings suggest that the divergent phase relationships observed in different experimental and computational studies of parkinsonian oscillations may stem partly from differences in the neuronal models employed, rather than solely from differences in connectivity architectures or external driving forces. This insight provides a new perspective on interpreting seemingly contradictory results in the literature and highlights the importance of carefully considering neuronal model selection when investigating oscillatory phenomena in the basal ganglia.

#### Results on the two-loop spiking network

Having established that different STN models (LIF vs. QIF) fundamentally determine the phase relationship between STN and Proto populations—with QIF neurons favoring in-phase locking and LIF neurons preferring anti-phase relationships—we now turn to the critical question of how these phase differences impact beta oscillation dynamics in the complete basal ganglia network. Our single-neuron and simplified network analyses provide the mechanistic foundation, but to fully test our hypothesis that contradictory findings in the literature stem from neuronal model choices rather than circuit-level disagreements, we must examine these phase relationships within the broader context of coupled subthalamopallidal and pallidostriatal loops. Thus, we performed comprehensive network simulations to demonstrate that the STN-Proto phase locking tendencies that we have characterized serve as the key determinant of whether STN enhances or suppresses pathological beta oscillations originating in the pallidostriatal system.

To investigate how single neuron dynamics influence beta oscillations in the full spiking network, we compared three network configurations: without STN (baseline), with LIF STN neurons, and with QIF STN neurons. Figure 11 shows the power spectra of Proto population firing rates for each condition. The baseline network without STN already generates beta oscillations through the pallidostriatal loop, serving as a benchmark for comparison. Remarkably, adding LIF STN neurons (red curve) actually decreases beta power compared to this baseline, while incorporating QIF STN neurons (green curve) substantially enhances beta oscillations, producing a more prominent peak around 15-18 Hz. This result demonstrates that the choice of single neuron model for STN critically determines whether the subthalamopallidal loop enhances or suppresses the beta oscillations generated by the pallidostriatal circuit, consistent with our rate model predictions that phase relationships between STN and Proto populations are key to oscillation amplitude.

**Fig 11.**
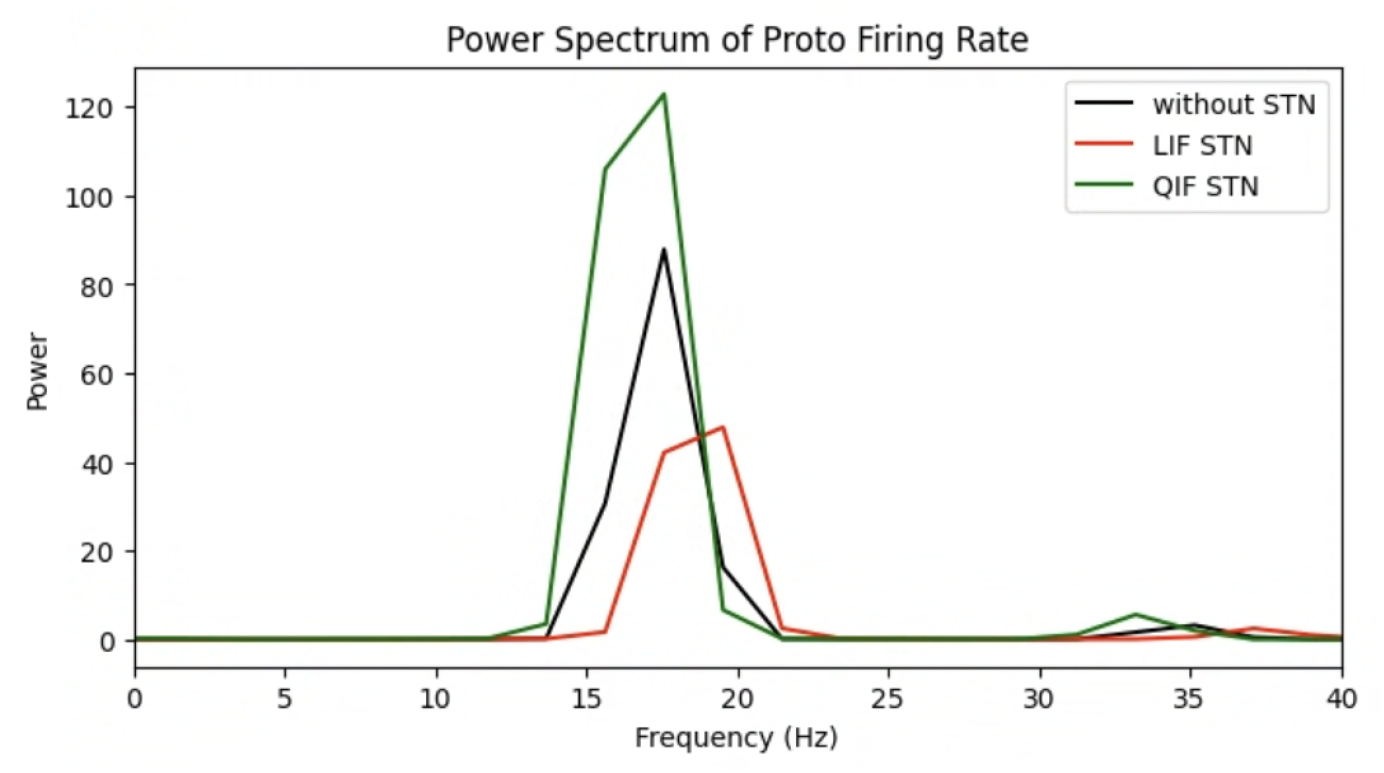
Power spectra of Proto population firing rates across network configurations. The baseline condition without STN (black) shows beta power generated by the pallidostriatal loop. Adding LIF STN neurons (red) reduces beta power compared to baseline, while QIF STN neurons (green) dramatically enhance beta oscillations with a more prominent peak around 15-18 Hz. Thus, single neuron dynamics in STN critically determine whether the subthalamopallidal loop enhances or suppresses beta oscillations generated by the pallidostriatal circuit.

To understand the mechanistic basis of how different STN neuron models affect beta oscillations, we performed activation experiments in the spiking networks. Figure 12 shows the effects of turning on the STN-to-Proto connection (green shaded region) for both LIF and QIF STN models. In the case of the LIF STN network (Fig 12A), activating the STN→Proto connection leads to a decrease in beta oscillation amplitude, as evidenced by both the blue shading in the difference spectrogram (top panel) and the comparison of the time series of Proto firing rates with (green) and without (orange) STN activation (middle panel). The bottom panel reveals that the phase difference between STN and Proto populations is 142.6 degrees during activation of the STN-to-Proto connection. In contrast, the QIF STN network (Fig 12B) shows the opposite effect: turning on the STN→Proto connection enhances beta oscillation amplitude and also affects the oscillation frequency slightly. The phase difference analysis (bottom panel) demonstrates that QIF STN establishes a markedly different phase relationship with Proto, with a phase difference of only 11.3 degrees. These results corroborate the idea that single neuron dynamics of STN determine the phase coupling with Proto, which in turn controls whether the subthalamopallidal loop enhances or suppresses beta oscillations generated by the pallidostriatal circuit.

**Fig 12.**
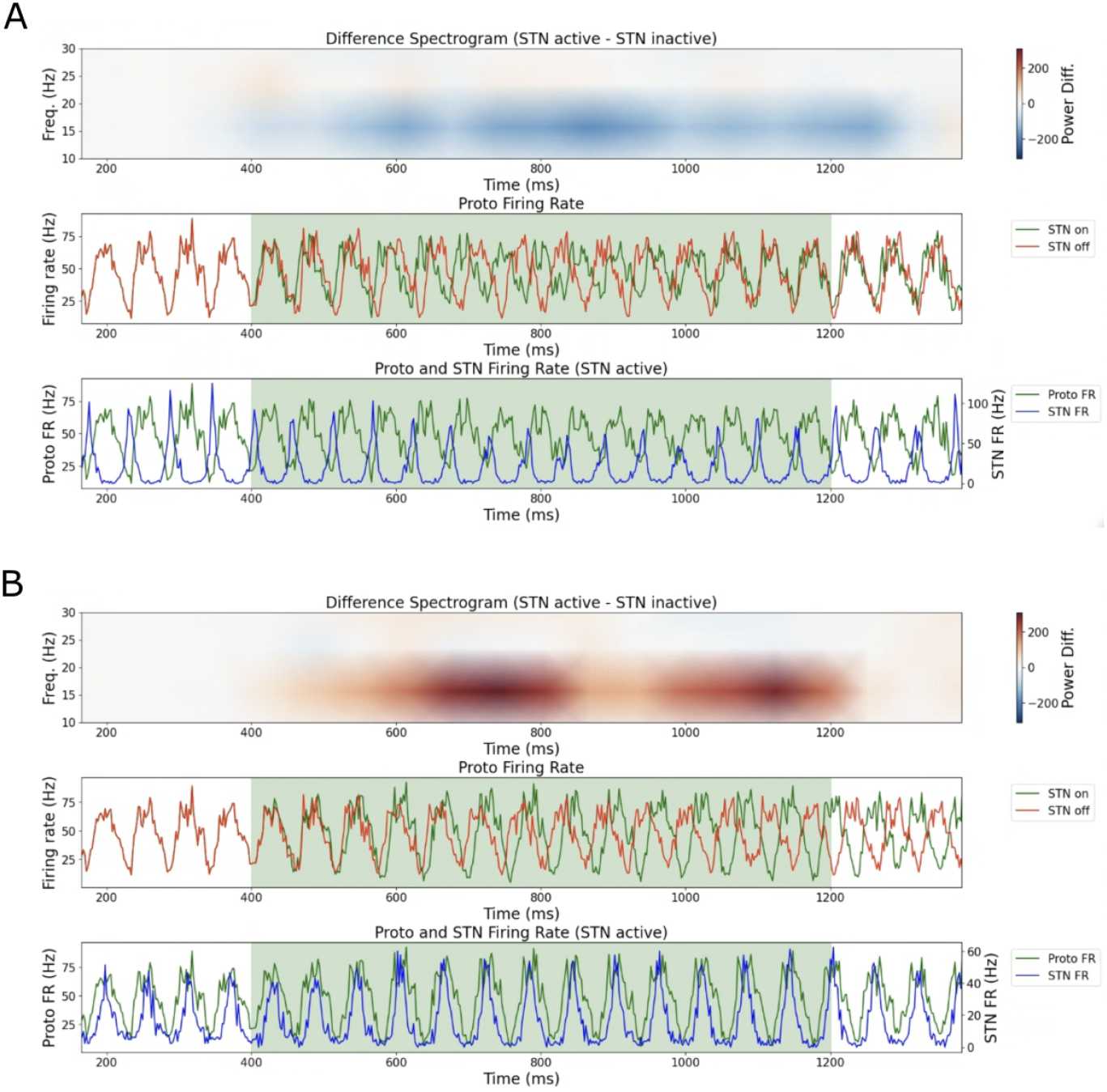
Effects of activation of STN→Proto connections on beta oscillations. (A) LIF STN configuration: turning on the STN→Proto connection (green region) decreases beta amplitude and establishes an anti-phase relationship between STN and Proto firing rates. (B) QIF STN configuration: the same intervention enhances beta amplitude and produces a different phase locking of STN and Proto.

**Fig 13.**
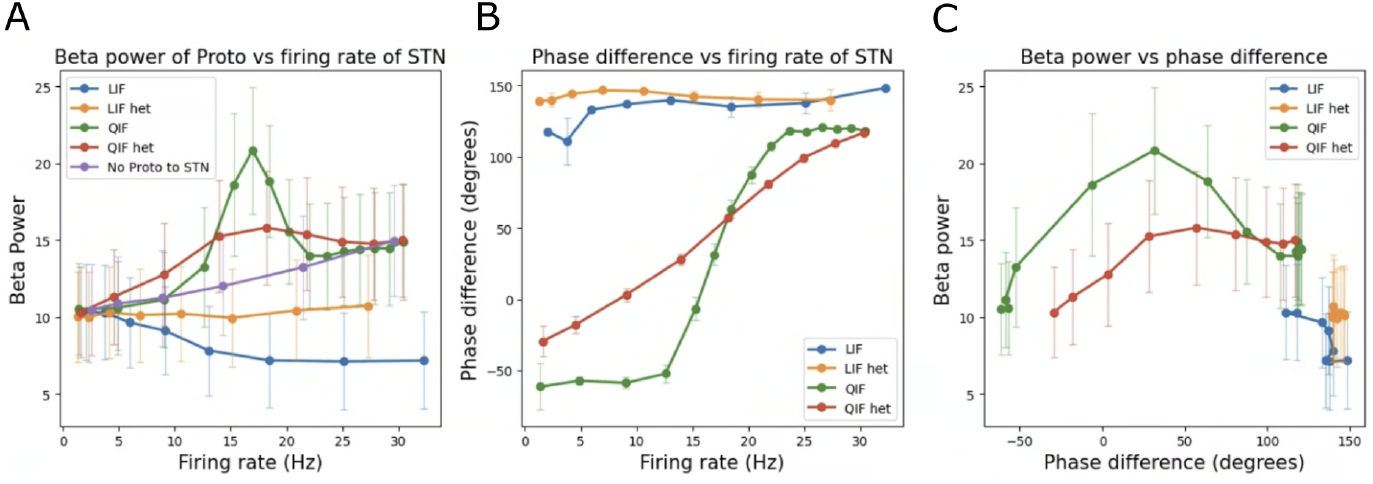
Effects of STN firing rate effects on beta power. (A) Beta power in Proto versus STN firing rate for different network configurations. The baseline case, without Proto→STN feedback (gray), shows an increase in beta with STN rate, while the inclusion of Proto→STN coupling reduces (LIF STN, red) or enhances (QIF STN, green) beta power. Heterogeneous input conditions (LIF het, QIF het) dampen the respective effects. (B) Phase difference between Proto and STN populations versus STN firing rate, showing that different neuron models achieve distinct phase relationships. (C) Beta power versus phase difference, demonstrating that phase coupling is the key mechanistic determinant of beta modulation. Higher phase differences reduce beta-enhancing effects regardless of neuron model. Error bars represent standard error across multiple seeds.

To systematically examine the relationship between STN firing rate, STN-Proto phase locking, and beta power modulation, we conducted parameter sweeps by varying the external input current to STN neurons across multiple network configurations. Figure 13 presents the results of this comprehensive analysis. Panel A shows how beta power in Proto varies with STN firing rate across various settings. The baseline condition without Proto→STN feedback (gray) demonstrates that increasing STN firing rate alone enhances beta power through direct excitatory effects on Proto. However, when Proto→STN feedback is present, the relationship becomes model-dependent: with LIF STN neurons (blue), beta power decreases compared to baseline, while with QIF STN neurons (green), it is enhanced. Notably, introducing heterogeneity in the input current of STN (LIF het and QIF het curves) partially dampens both the beta-reducing and beta-enhancing effects of the respective neuron models. Panel B hints at the mechanistic basis for the QIF results by showing how phase difference between Proto and STN varies with STN firing rate. In the QIF case without heterogeneity, the phase difference between STN and Proto rises sharply through the in-phase range, leading to a strong enhancement of beta power over a relatively narrow range of STN firing rates. Heterogeneity smooths out the phase difference curve, yielding a milder beta enhancement but over a broader range of STN rates. Panel C directly demonstrates this principle by plotting beta power against phase difference, revealing that higher phase differences reduce the beta-enhancing effect regardless of the underlying neuron model. Even for QIF STN, which generally enhances beta oscillations, larger phase differences diminish this amplifying effect.

## Discussion

Our investigation into how neuronal model choice affects beta oscillation dynamics in the basal ganglia has revealed a fundamental mechanistic principle that reconciles seemingly contradictory findings in the Parkinson’s disease literature. Through systematic analysis spanning rate models, single neuron analysis, and spiking network simulations, we demonstrated that the phase relationship between STN and Proto populations serves as the critical determinant of whether the subthalamopallidal loop enhances or suppresses beta oscillations originating in the pallidostriatal circuit. Quadratic integrate-and-fire (QIF) STN neurons demonstrate stable in-phase relationships with inhibitory input, while leaky integrate-and-fire (LIF) STN neurons exhibit anti-phase locking. These neuron-level distinctions translate to population-level dynamics, determining whether the subthalamopallidal loop acts as a beta amplifier or suppressor.

Our rate model and spiking network simulations provided converging evidence that phase relationships directly control oscillation amplitude, with experiments demonstrating that identical circuit architectures produce opposite effects on beta power depending solely on STN neuron model choice. Parameter sweeps confirmed that these phase preferences remain consistent across physiologically relevant firing rate ranges, even when neurons are not strictly phase-locked to network rhythms.

The finding that QIF STN neurons establish in-phase relationships with Proto while LIF STN neurons prefer anti-phase coupling provides a direct mechanistic explanation for the divergent results between [39] and [40]. Rather than representing fundamental disagreements about circuit mechanisms, these studies’ contradictory conclusions likely stemmed from different neuronal model implementations. This observation underscores the point that careful consideration of neuronal model selection is crucial when investigating oscillatory phenomena in neural circuits.

Because STN neurons excite their Proto counterparts, an in-phase relationship in their firing tends to strengthen ongoing Proto oscillations, and this effect manifests with QIF STN neurons. Alternatively, with anti-phase locking, STN excitation arrives at troughs in Proto firing, diminishing these minima, and wanes at peaks in Proto firing, removing any potential boosting effect. Thus, anti-phase STN-Proto relationships lead to suppression of Proto oscillations, as we have seen with LIF STN neurons. In the LIF case, input heterogeneity dampens oscillations in STN firing, thereby weakening the impact of anti-phase STN firing on Proto beta (Fig 10).

The link between a particular STN model *v*^*′*^ = *I* + *f* (*v*) −*I*_*inh*_ and the STN-Proto phase relationship that it produces derives from the shape of the curve *I* + *f* (*v*) = 0. This curve determines where and when *dv/dt* slows down and how this deceleration is affected by inhibition, which combine to set the range of input phases over which a spike can occur (Fig 6). We can expect that qualitatively similar ideas will arise in higher-dimensional neuronal models, although we decided to exclude these from this study both to stay true to the original works on this topic [39, 40] and to avoid drowning in degrees of freedom and parameter choices that could obscure the fundamental effects that we have described.

The time-to-spike (TTS) framework developed here represents a significant extension of previous applications [45, 46], demonstrating that TTS can be redefined as a map based on comparison with a reference signal instead of based directly on time elapsed since the last spike as was done previously. By redefining TTS based on temporal offset from periodic inhibition, we captured phase dynamics while reducing complex continuous dynamics to mathematically tractable discrete maps. This approach is particularly well-suited for excitatory-inhibitory networks in which sequential interactions provide natural discrete events for mapping.

An important consideration emerging from our literature review is the potential for species-specific differences in basal ganglia dynamics. Experiments supporting STN’s role in beta generation [17, 18] were conducted in primates, while studies showing limited STN effects on beta [27] and supporting pallidostriatal origins of beta oscillations [5, 28–34] used non-primates. This pattern suggests that there may be differences in STN neuron excitability between species, which could explain apparent contradictions through the phase relationship mechanisms we identified. Direct measurement of phase relationships between STN and Proto populations during beta oscillations in both primate and rodent models could be used to test for the presence of the relationships that we predict. Additionally, applying periodic inhibitory stimulation to single STN neurons in slice preparations could directly test whether primate and rodent STN neurons exhibit QIF-like and LIF-like dynamics, respectively, at least with regard to their phase-locking properties. Testing periodic inhibition at dendritic versus somatic locations could reveal whether dendritic integration, potentially more extensive in primate STN neurons, contributes to QIF-like behavior.

While computational efficiency often drives the use of simplified integrate-and-fire models, our findings demonstrate that this choice fundamentally alters biological conclusions. The field would benefit from systematic comparisons between simplified and conductance-based models, specifically examining whether they preserve critical dynamical properties like phase relationships. The demonstration that single neuron dynamics can fundamentally alter population-level pathological activity represents a broader principle that should be considered from both theoretical and experimental perspectives. On the computational side, seemingly minor modeling choices can have profound consequences for model network behavior and clinical interpretation [6]. This insight underscores the importance of validating computational predictions against multiple experimental measures and considering how model assumptions might influence conclusions about disease mechanisms and therapeutic targets. On the biological side, these results offer a reminder that even within large networks of coupled neurons, the properties of individual neurons can influence emergent dynamics and remain important to elucidate.

Beyond Parkinson’s disease, our work provides insights into how pathological oscillations may emerge and be modulated in other neurological and psychiatric conditions. The principle that phase relationships between brain regions critically determine oscillatory power could apply to epilepsy, schizophrenia, and other disorders characterized by abnormal neural synchronization [47–50].

## Supporting information

### S1 Appendix – Proof that the LIF STN model fires during the inhibition-off period

We study the existence and stability of fixed points of the time-to-spike map arising in the inhibition-on and -off periods for an LIF neuron driven with periodic inhibition. For convenience, we first obtain the solution of the LIF equation for a general input 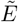. The equation is given by 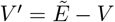, and with data *V* (*t*_0_) = *V*_0_ the solution is

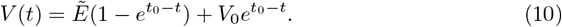

Let *I* and *a* be the input current and inhibition strength, respectively. Then 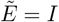 when inhibition is off and 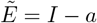 when inhibition is on. Suppose that when *V* reaches the threshold at *V* = *V*_*th*_ = 1 it is reset to *V* = 0 and that we start the solution with *V* (0) = 0, with the neuron having just been reset after a spike.

We will proceed by showing that the TTS map acts as a contraction everywhere on the interval of initial conditions corresponding to spiking and being reset with inhibition off, *t*^*in*^ ∈ (*T/*4, 3*T/*4). This result will imply that any fixed point arising in the time range corresponding to inhibition off will be stable. Recall from section, as visualized by Figure 5, that *t*^*in*^ is defined as the time difference between the spike of the excitatory neuron and the mid-point of the next time interval when its inhibitory input is on. In this case, the first positive time when inhibition turns on will be at *t* = *t*^*in*^ −*T/*4. The next inhibition-off time will be *t*^*in*^ −*T/*4 + *T/*2 = *t*^*in*^ + *T/*4 and inhibition will remain off up to *t*^*in*^ + 3*T/*4. The full time domain of interest in this case is [0, *t*^*in*^ + 3*T/*4], because we want the neuron fire again during this inhibition-off period. Thus, the relevant piecewise system takes the form

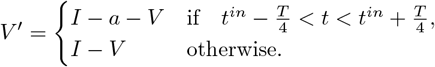

Consider the moments when inhibition turns on 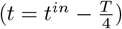 and off 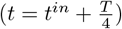, respectively. The solution formula (10) gives us

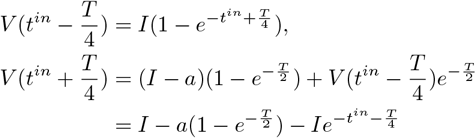

Since we are assuming that firing occurs in the inhibition-off period, the voltage reaches the firing threshold after time 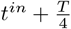, say at 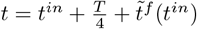. Applying equation (10) at the moment the neuron spikes with 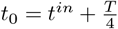 and grouping the exponential terms gives us

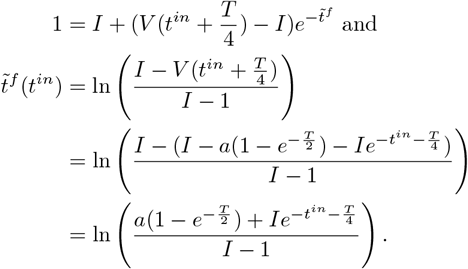

Now, suppose that *t*^*in*,1^ *< t*^*in*,2^ are two possible firing times during the inhibition-off period. Since 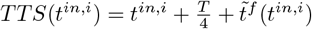,

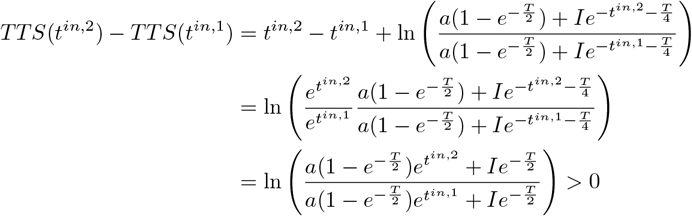

and, using the time-to-spike map definition (4),

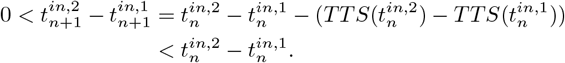

Thus, we have shown that the map is a contraction on the inhibition-off period, and hence any fixed point within that period is stable.

For such a fixed point to exist, we need

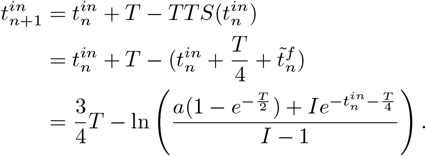

Thus, the fixed point equation becomes

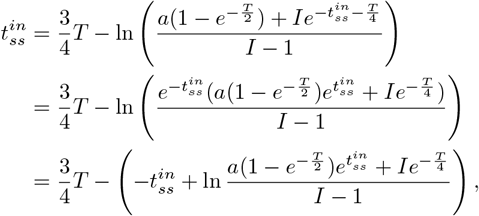

from which we derive

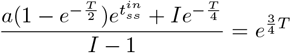

and hence

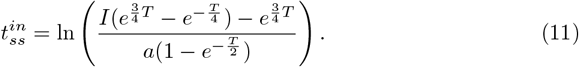

So, the fixed point exists, and is an increasing function of *I* and a decreasing function of *a*, if this expression is well-defined and lies in the interval (*T/*4, 3*T/*4), corresponding to the inhibition-off period. We are only interested in the case when *I >* 1, so that the neuron can fire, and the conditions 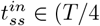, 3*T/*4) are equivalent to the conditions

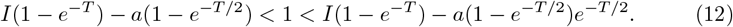

Thus, the LIF model gives a stable fixed point with inhibition-off firing when conditions (12) hold or, rewritten in terms of conditions on *I* or *a*,

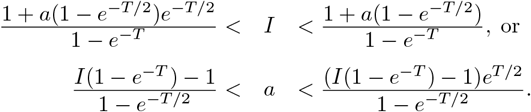

To complete our argument, we want to show that the map acts locally as a dilation on the region of initial conditions with inhibition on. This result will imply that any fixed point in the inhibition-on region is unstable. To proceed, assume that the neuron spiked with inhibition on. Then, the next inhibition-off time would be 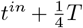, and the next inhbition on time would be 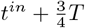; thus,

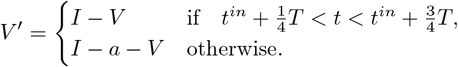

Following the steps in the last case, we have:

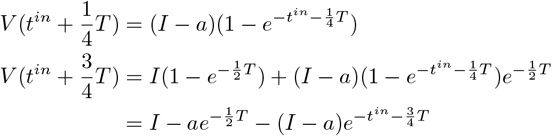

and

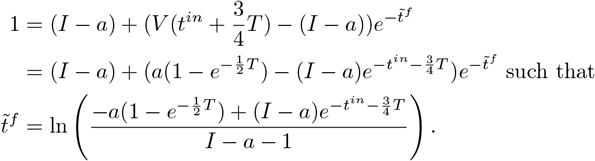

Proceeding as in the previous case and using 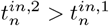, we have

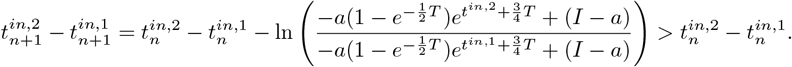

Hence, the map is a dilation on initial conditions in the inhibition-on region and cannot have a stable fixed point there.

### S2 Appendix – Proof that the QIF fires during the inhibition-on period

The goal of this section is to show that for the QIF model, there exists a fixed point of the time-to-spike map inside the inhibition-on interval and to derive conditions under which this fixed point is stable. Similar to what we did in the LIF case, we compute the solution of the QIF equation with an arbitrary input 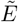.

### Case 1: The nullcine stays above the *x*-axis

We first consider the case *I >* 0, *a >* 0, and *I* − *a >* 0, so that the *V*-nullcline does not intersect the axis when inhibition is on. If we ignore the threshold and associated reset, for 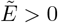, we have:

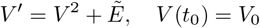

with solution

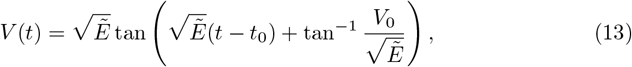

which we can invert to obtain

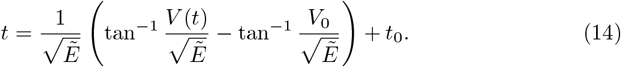

Again, 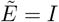 when inhibition is off and 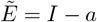 when inhibition is on. Suppose that the model neuron fires with inhibition on. The voltage ODE becomes

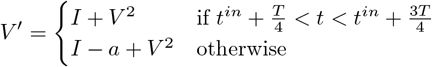

where *t*^*in*^ ∈ ( −*T/*4, *T/*4) is positive if the firing happens before the midpoint of the inhibition-on interval and negative if it happens afterwards.

Consider the voltages the moments when inihibition turns off 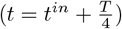 and on 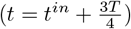, respectively. The solution formula (13) gives

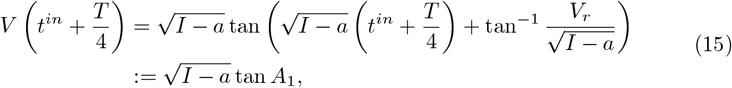

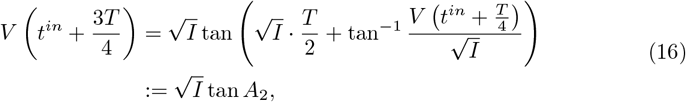

where *A*_1_ and *A*_2_ will be used for the stability argument below.

### Existence

Consider *t*^*in*^ as a parameter, the solution formula (13) applied from the time *t* = *t*^*in*^ + 3*T/*4 when inhibition turns back on up to time *T* gives

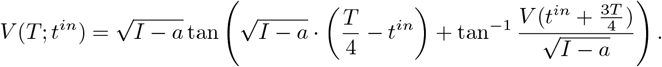

To establish the existence of a fixed point, we want to show that the trajectory with 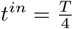 does not reach the firing threshold by the time inhibition turns on again at *t* = *T*, while the trajectory with 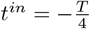, ends up past the firing threshold at *t* = *T* (ignoring reset). In other words, we want

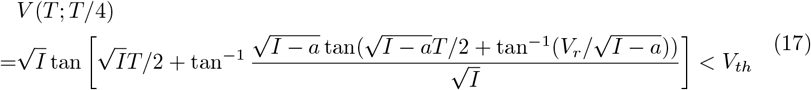

and

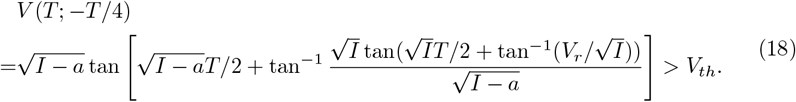

If conditions (17) and (18) are satisfied, then by the intermediate value theorem, there exists 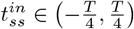 such that

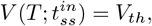

with reset back to 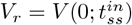, so the corresponding solution is phase-locked to the periodic inhibition with firing during the inhibition-on period; that is, this solution is our desired fixed point.

Intuitively, we expect it to be easy to satisfy both (17) and (18): If the inhibition-on period comes first, as for (17), then the trajectory cannot advance much in *V* as long as *I* − *a* is relatively small (i.e., due to the ghost of the saddle node bifurcation at *I* − *a* = 0), and then it has a long way to go to reach threshold once inhibition is off. On the other hand, if inhibition-off comes first, as for (18), then the trajectory can freely advance past what would have been a bottleneck had inhibition been present, and then it is relatively free to progress to threshold once inhibition comes on.

To demonstrate this intuition more analytically, let’s first consider the solution that has *t*^*in*^ = *T/*4, which starts from *t* = 0 with *V* (0) = *V*_*r*_ *<* 0. Without loss of generality, let *V*_*th*_ = 1. After time *T*, this solution’s voltage value is given by the left hand side of inequality (17). Suppose we approximate *V* (*T/*2), when inhibition turns off, by 0. Then 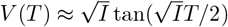. Next, consider the solution 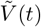 that has *t*^*in*^ = −*T/*4, with 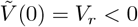. This solution has 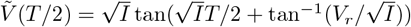. To find 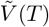, we integrate 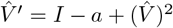 with 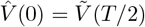, up to time *T/*2. Assuming that *I* − *a* is small, we approximate 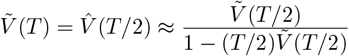, from the solution of 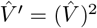.

Thus, we ask, under what conditions will we have *V* (*T*) *<* 1, so that (17) holds, and 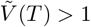, so that (18) holds? Using our approximation, the latter condition is equivalent to 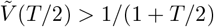. Note that 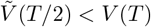, with 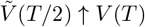 as *V*_*r*_ ↑ 0. Thus, since 1*>* 1*/*(1+ *T/*2), for any fixed *T*, we can always satisfy these conditions by choosing 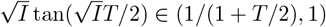 and *V*_*r*_ not too far below 0.

### Stability

We show that any fixed point in the inhibition-on region is stable by calculat ing the de rivative of the map *TTS*. Equation (14) with 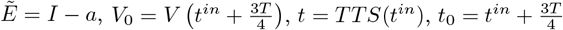, and *V* (*TTS*(*t*^*in*^)) = *V*_*th*_ yields

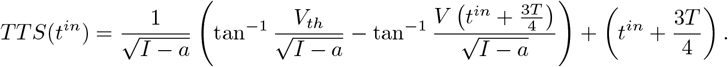

We apply chain rule to compute the derivative of TTS:

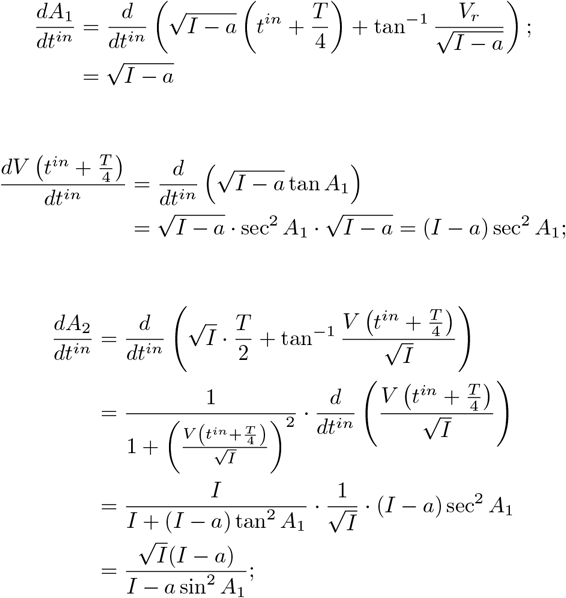

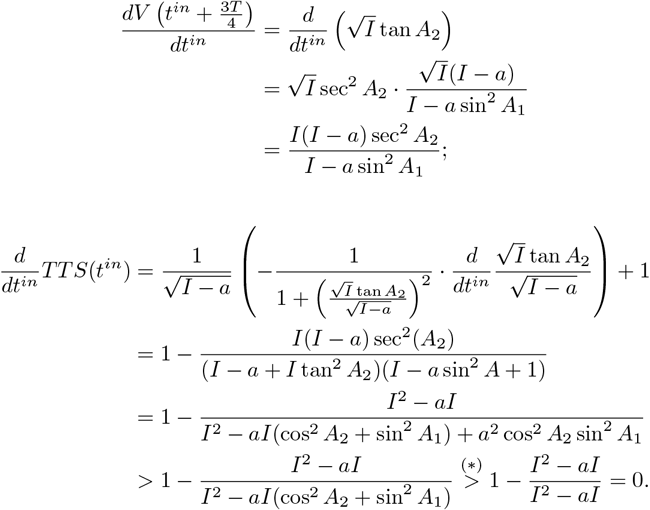

Inequality (*) would be true if the following conditions are satisfied:

1. *A*_1_, *A*_2_ *>* 0

2. 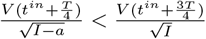

That is, under these conditions, by definitions (15) and (16), we have

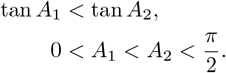

Since cos^2^(*x*) is a decreasing function in 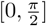, it follows that

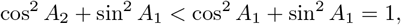

which establishes inequality (*).

Note that *A*_2_ *>* 0 from condition (a) is equivlent to *V* (*t*^*in*^ + 3*T/*4) *>* 0 by definition (16). This is usually satisfied in practice, because when inhibition turns on, the voltage should be close enough to the firing threshold, and we always set the threshold to be positive. Similarly, condition (b) means that the voltage during the inhibition-off period should increase sufficiently. The condition *A*_1_ *>* 0 could be replaced by *A*_2_ + *A*_1_ *>* 0 with *A*_1_ ∈ ( −*π/*2, 0]. This condition certainly holds if *V*_*r*_ is not too far below 0.

Since *I > a*, the fractional term in 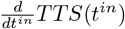 is positive, which implies 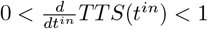. Therefore, the fixed point of the map 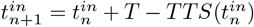 during inhibition is stable under conditions (a) and (b), as the derivative of the map satisfies |*f* ^*′*^| *<* 1.

### Case 2: The nullcine intersects the axis

We now assume *I >* 0, *a >* 0, and *I* − *a <* 0. As in the previous case, we have

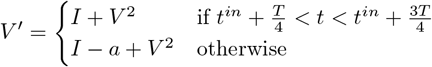

We will also assume here that 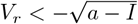, such that *V* ^*′*^ *>* 0 immediately after any reset regardless of its timing. Note that if inhibition turns on, giving 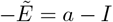, with *t* = *t*_0_, *V* (*t*_0_) = *V*_0_, and 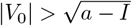 then 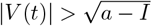 remains true as long as inhibition stays on.

The general solution to the QIF equation 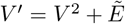 with constant input 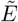 is given by equation (13) when 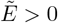. When 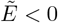 and 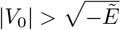, the solution is

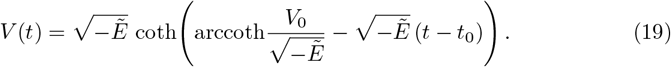

We now use these solutions to compute the time-to-spike function. Using (19) on the inhibition-on segment with 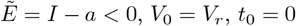, and evaluating at 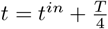 gives:

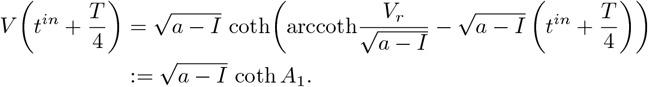

Using (13) on the inhibition-off segment with 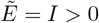 over a time duration *T/*2 starting from *t* = *t*^*in*^ + *T/*4 yields:

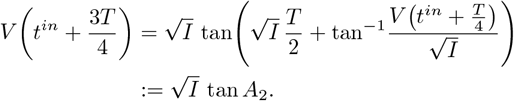

### Existence

We will derive conditions for the existence of a fixed point under the assumption that 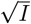 tan 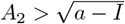. Considering *t*^*in*^ as a parameter, the solution formula (19) applied from the time *t* = *t*^*in*^ + 3*T/*4 when inhibition turns back on up to time *T* thus gives

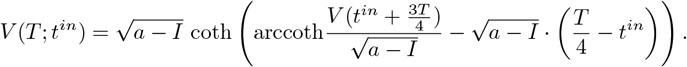

To establish the existence of a fixed point, we want to show that the trajectory with 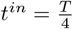 does not reach the firing threshold by the time inhibition turns on again at *t* = *T*, while the trajectory with 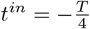, ends up past the firing threshold at *t* = *T* (ignoring reset). In other words, we want

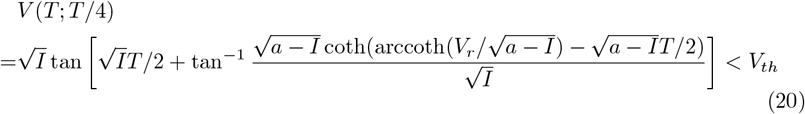

and

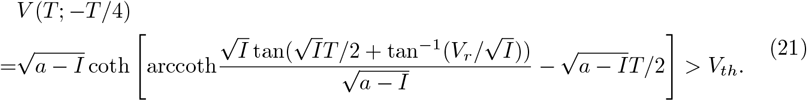

Under our earlier assumption that 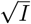 tan 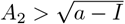, if conditions (20) and (21) are satisfied, then by the intermediate value theorem, there exists 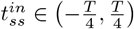, such that

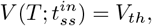

with reset back to 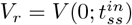, so the corresponding solution is phase-locked to the periodic inhibition with firing during the inhibition-on period; that is, this solution is our desired fixed point.

### Stablity

The time-to-spike function satisfies *V* (TTS(*t*^*in*^)) = *V*_th_. Similarly to the previous case, we use (19) with *t*_0_ = *t*^*in*^ + 3*T/*4, *V*_0_ = *V* (*t*^*in*^ + 3*T/*4), and 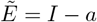 to calculate that when spiking occurs during the inhibition-on period,

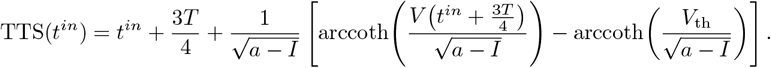

To compute the derivative of TTS with respect to *t*^*in*^, we first compute several intermediate derivatives. For brevity, define 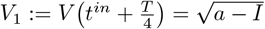 coth *A*_1_ and 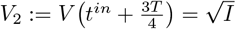 tan *A*_2_. Then

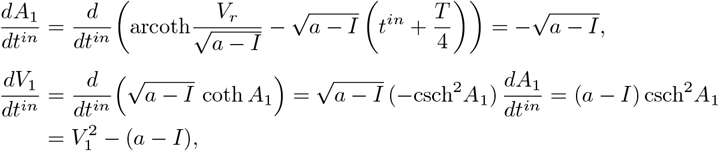

where we used the identity coth^2^ *A*_1_ − csch^2^*A*_1_ = 1.

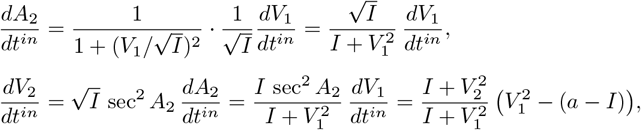

using the identity sec^2^ *A*_2_ = 1 + tan^2^ *A*_2_ = 1 + *V* ^2^*/I*.

Finally, differentiating TTS with respect to *t*^*in*^ gives

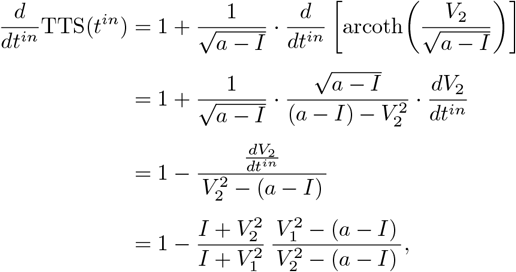

where we used 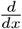 arcoth 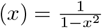. For a fixed point 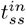 of the map to be stable, we require 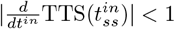. Recall that 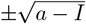 are the fixed points of the ode with 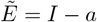. Since we are assuming that the neuron fires while inhibition is on, the voltage must be blocked by the stable fixed point when inhibition is on at the start of the cycle 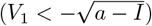 and must have already exceeded the unstable fixed point when inhibition turns on again later in the cycle 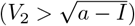. Hence, the fractional term is always positive, and we have 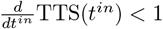. The condition 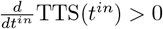 is equivalent to

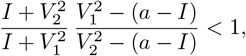

which simplifies to

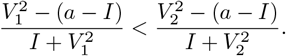

Since the function 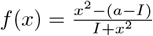 is monotonically increasing for *x >* 0, this condition is satisfied if and only if *V*_1_ *< V*_2_. During the inhibition-off period from *t*^*in*^ + *T/*4 to *t*^*in*^ + 3*T/*4, the neuron evolves under *V* ^*′*^ = *I* + *V* ^2^ with *I >* 0, so *V* ^*′*^ *>* 0 and the voltage is strictly increasing. Therefore, *V*_1_ *< V*_2_ is automatically satisfied, ensuring that the inhibition-off fixed point is stable.

## Acknowledgments

The authors received partial support from NIH awards R01NS125814 (KNT, JER) and R01DA059993 (JER).

## References

1. Hutchison WD, Dostrovsky JO, Walters JR, Courtemanche R, Boraud T, Goldberg J, et al. Neuronal oscillations in the basal ganglia and movement disorders: evidence from whole animal and human recordings. Journal of Neuroscience. 2004;24(42):9240–9243. doi:10.1523/JNEUROSCI.3366-04.2004.

2. Yager LM, Garcia AF, Wunsch AM, Ferguson SM. The ins and outs of the striatum: role in drug addiction. Neuroscience. 2015;301:529–541.

3. Koós T, Tepper JM. Inhibitory control of neostriatal projection neurons by GABAergic interneurons. Nature Neuroscience. 1999;2(5):467–472.

4. Parent A, Hazrati LN. Functional anatomy of the basal ganglia. I. The cortico-basal ganglia-thalamo-cortical loop. Brain research reviews. 1995;20(1):91–127.

5. Mallet N, Micklem BR, Henny P, Brown MT, Williams C, Bolam JP, et al. Dichotomous Organization of the External Globus Pallidus. Neuron. 2012;74(6):1075–1086. doi:10.1016/j.neuron.2012.04.027.

6. Rubin JE. Computational models of basal ganglia dysfunction: the dynamics is in the details. Current opinion in neurobiology. 2017;46:127–135.

7. Mirzaei A, Kumar A, Leventhal D, Mallet N, Aertsen A, Berke J, et al. Sensorimotor processing in the basal ganglia leads to transient beta oscillations during behavior. Journal of Neuroscience. 2017;37(46):11220–11232.

8. Bergman H, Wichmann T, DeLong MR. Reversal of experimental parkinsonism by lesions of the subthalamic nucleus. Science. 1990;249(4975):1436–1438.

9. Bergman H, Wichmann T, Karmon B, DeLong MR. The primate subthalamic nucleus. II. Neuronal activity in the MPTP model of parkinsonism. Journal of Neurophysiology. 1994;72(2):507–520. doi:10.1152/jn.1994.72.2.507.

10. Kühn AA, Kupsch A, Schneider GH, Brown P. Reduction in subthalamic 8-35 Hz oscillatory activity correlates with clinical improvement in Parkinson’s disease. European Journal of Neuroscience. 2006;23(7):1956–1960. doi:10.1111/j.1460-9568.2006.04717.x.

11. Alavi SM, Mirzaei A, Valizadeh A, Ebrahimpour R. Excitatory deep brain stimulation quenches beta oscillations arising in a computational model of the subthalamo-pallidal loop. Scientific Reports. 2022;12(1):1–20. doi:10.1038/s41598-022-07540-2.

12. Hammond C, Bergman H, Brown P. Pathological synchronization in Parkinson’s disease: networks, models and treatments. Trends in Neurosciences. 2007;30(7):357–364. doi:10.1016/j.tins.2007.05.004.

13. Vitek JL, Zhang J, Hashimoto T, Russo GS, Baker KB. External pallidal stimulation improves parkinsonian motor signs and modulates neuronal activity throughout the basal ganglia thalamic network. Experimental Neurology. 2012;233(1):581–586. doi:10.1016/j.expneurol.2011.09.031.

14. Deffains M, Iskhakova L, Katabi S, Israel Z, Bergman H. Longer β Oscillatory Episodes Reliably Identify Pathological Subthalamic Activity in Parkinsonism. Movement Disorders. 2018;33(10):1609–1618.

15. Yu Y, Sanabria DE, Wang J, Hendrix CM, Zhang J, Nebeck SD, et al. Parkinsonism Alters Beta Burst Dynamics Across the Basal Ganglia–Motor Cortical Network. Journal of Neuroscience. 2021;41(10):2274–2286.

16. Plenz D, Kital ST. A basal ganglia pacemaker formed by the subthalamic nucleus and external globus pallidus. Nature. 1999;400(6745):677–682. doi:10.1038/23281.

17. Deffains M, Iskhakova L, Katabi S, Haber SN, Israel Z, Bergman H. Subthalamic, not striatal, activity correlates with basal ganglia downstream activity in normal and parkinsonian monkeys. Elife. 2016;5:e16443.

18. Tachibana Y, Iwamuro H, Kita H, Takada M, Nambu A. Subthalamo-pallidal interactions underlying parkinsonian neuronal oscillations in the primate basal ganglia. European Journal of Neuroscience. 2011;34(9):1470–1484.

19. Wilson HR, Cowan JD. Excitatory and inhibitory interactions in localized populations of model neurons. Biophysical journal. 1972;12(1):1–24.

20. Terman D, Rubin JE, Yew A, Wilson C. Activity patterns in a model for the subthalamopallidal network of the basal ganglia. Journal of Neuroscience. 2002;22(7):2963–2976.

21. Gillies, Willshaw D, Li Z. Subthalamic–pallidal interactions are critical in determining normal and abnormal functioning of the basal ganglia. Proceedings of the Royal Society of London Series B: Biological Sciences. 2002;269(1491):545–551.

22. Holgado AJN, Terry JR, Bogacz R. Conditions for the generation of beta oscillations in the subthalamic nucleus–globus pallidus network. Journal of Neuroscience. 2010;30(37):12340–12352.

23. Kumar A, Cardanobile S, Rotter S, Aertsen A. The role of inhibition in generating and controlling Parkinson’s disease oscillations in the basal ganglia. Frontiers in systems neuroscience. 2011;5:86.

24. Pasillas-Lépine W. Delay-induced oscillations in Wilson and Cowan’s model: an analysis of the subthalamo-pallidal feedback loop in healthy and parkinsonian subjects. Biological cybernetics. 2013;107(3):289–308.

25. Bahuguna J, Sahasranamam A, Kumar A. Uncoupling the roles of firing rates and spike bursts in shaping the STN-GPe beta band oscillations. PLoS Computational Biology. 2020;16(3):e1007748.

26. Barraza D, Kita H, Wilson CJ. Slow spike frequency adaptation in neurons of the rat subthalamic nucleus. Journal of neurophysiology. 2009;102(6):3689–3697.

27. Crompe Bdl, Aristieta A, Leblois A, Elsherbiny S, Boraud T, Mallet NP. The globus pallidus orchestrates abnormal network dynamics in a model of Parkinsonism. Nature communications. 2020;11(1):1570.

28. Abdi A, Mallet N, Mohamed FY, Sharott A, Dodson PD, Nakamura KC, et al. Prototypic and arkypallidal neurons in the dopamine-intact external globus pallidus. Journal of Neuroscience. 2015;35(17):6667–6688. doi:10.1523/JNEUROSCI.4662-14.2015.

29. Bevan MD, Booth PA, Eaton SA, Bolam JP. Selective innervation of neostriatal interneurons by a subclass of neuron in the globus pallidus of the rat. Journal of Neuroscience. 1998;18(22):9438–9452. doi:10.1523/JNEUROSCI.18-22-09438.1998.

30. Fujiyama F, Nakano T, Matsuda W, Furuta T, Udagawa J, Kaneko T. A single-neuron tracing study of arkypallidal and prototypic neurons in healthy rats. Brain Structure and Function. 2016;221(9):4733–4740. doi:10.1007/s00429-015-1152-2.

31. Cagnan H, Mallet N, Moll CK, Gulberti A, Holt AB, Westphal M, et al. Temporal evolution of beta bursts in the parkinsonian cortical and basal ganglia network. Proceedings of the National Academy of Sciences. 2019;116(32):16095–16104. doi:10.1073/pnas.1819975116.

32. Dodson PD, Larvin JT, Duffell JM, Garas FN, Doig NM, Kessaris N, et al. Distinct developmental origins manifest in the specialized encoding of movement by adult neurons of the external globus pallidus. Neuron. 2015;86(2):501–513. doi:10.1016/j.neuron.2015.03.040.

33. Hernández VM, Hegeman DJ, Cui Q, Kelver DA, Fiske MP, Glajch KE, et al. Parvalbumin+ neurons and Npas1+ neurons are distinct neuron classes in the mouse external globus pallidus. Journal of Neuroscience. 2015;35(34):11830–11847. doi:10.1523/JNEUROSCI.0672-15.2015.

34. Mastro KJ, Bouchard RS, Holt HAK, Gittis AH. Transgenic mouse lines subdivide external segment of the globus pallidus (GPe) neurons and reveal distinct GPe output pathways. Journal of Neuroscience. 2014;34(6):2087–2099. doi:10.1523/JNEUROSCI.4646-13.2014.

35. Corbit VL, Whalen TC, Zitelli KT, Crilly SY, Rubin JE, Gittis AH. Pallidostriatal projections promote β oscillations in a dopamine-depleted biophysical network model. Journal of Neuroscience. 2016;36(20):5556–5571.

36. Leblois A, Boraud T, Meissner W, Bergman H, Hansel D. Competition between feedback loops underlies normal and pathological dynamics in the basal ganglia. Journal of Neuroscience. 2006;26(13):3567–3583.

37. McCarthy M, Moore-Kochlacs C, Gu X, Boyden E, Han X, Kopell N. Striatal origin of the pathologic beta oscillations in Parkinson’s disease. Proceedings of the national academy of sciences. 2011;108(28):11620–11625.

38. Damodaran S, Cressman JR, Jedrzejewski-Szmek Z, Blackwell KT. Desynchronization of fast-spiking interneurons reduces β-band oscillations and imbalance in firing in the dopamine-depleted striatum. Journal of Neuroscience. 2015;35(3):1149–1159.

39. Ortone A, Vergani AA, Ahmadipour M, Mannella R, Mazzoni A. Dopamine depletion leads to pathological synchronization of distinct basal ganglia loops in the beta band. PLoS computational biology. 2023;19(4):e1010645.

40. Lindi SA, Mallet NP, Leblois A. Synaptic changes in pallidostriatal circuits observed in the parkinsonian model triggers abnormal beta synchrony with accurate spatio-temporal properties across the basal ganglia. Journal of Neuroscience. 2024;44(9).

41. Mallet N, Pogosyan A, Márton LF, Bolam JP, Brown P, Magill PJ. Parkinsonian beta oscillations in the external globus pallidus and their relationship with subthalamic nucleus activity. Journal of neuroscience. 2008;28(52):14245–14258.

42. Gage GJ, Stoetzner CR, Wiltschko AB, Berke JD. Selective activation of striatal fast-spiking interneurons during choice execution. Neuron. 2010;67(3):466–479.

43. Berke JD, Okatan M, Skurski J, Eichenbaum HB. Oscillatory entrainment of striatal neurons in freely moving rats. Neuron. 2004;43(6):883–896.

44. Miller BR, Walker AG, Shah AS, Barton SJ, Rebec GV. Dysregulated information processing by medium spiny neurons in striatum of freely behaving mouse models of Huntington’s disease. Journal of neurophysiology. 2008;100(4):2205–2216.

45. Tse KN, Ermentrout GB, Rubin JE. A Nonlinear Map Describing Relaxation to Cluster States in an Adapting Neuronal Network. SIAM Journal on Applied Dynamical Systems. 2025;24(2):1585–1621.

46. Krupa M, Gielen S, Gutkin B. Adaptation and Shunting Inhibition Leads to Pyramidal/Interneuron Gamma with Sparse Firing of Pyramidal Cells. Journal of Computational Neuroscience. 2014;37(2):357–376.

47. Nimmrich V, Draguhn A, Axmacher N. Neuronal network oscillations in neurodegenerative diseases. Neuromolecular medicine. 2015;17(3):270–284.

48. Jiruska P, Alvarado-Rojas C, Schevon CA, Staba R, Stacey W, Wendling F, et al. Update on the mechanisms and roles of high-frequency oscillations in seizures and epileptic disorders. Epilepsia. 2017;58(8):1330–1339.

49. Singh A. Oscillatory activity in the cortico-basal ganglia-thalamic neural circuits in Parkinson’s disease. European Journal of Neuroscience. 2018;48(8):2869–2878.

50. Hirano Y, Uhlhaas PJ. Current findings and perspectives on aberrant neural oscillations in schizophrenia. Psychiatry and Clinical Neurosciences. 2021;75(12):358–368.

